# Human XIST RNA acts early to condense architecture which facilitates A-repeat density-dependent initiation of gene silencing

**DOI:** 10.1101/2022.02.04.479153

**Authors:** Melvys Valledor, Meg Byron, Brett Dumas, Dawn M. Carone, Lisa L. Hall, Jeanne B. Lawrence

## Abstract

XIST RNA triggers gene silencing chromosome-wide and transforms a euchromatic chromosome into a condensed Barr body. XIST is heavily studied in mouse ES cells, but here an inducible iPSC system allows analysis of initial steps in *human* chromosome silencing, revealing key points not known in either system. XIST RNA distribution was examined relative to biochemical and transcriptional changes directly within architecture of individual chromosome territories. Within a few hours of induction, XIST transcripts distribute as a large “sparse zone” and a smaller “dense zone”, which, importantly, exhibit different effects on chromatin. Very sparse transcripts immediately trigger bright staining for H2AK119ub and CIZ1, a structural matrix protein. In contrast, H3K27me3 enrichment comes hours later and is much more restricted to the smaller dense RNA zone, which enlarges as the chromosome condenses. Importantly, silencing of several genes examined occurred well after architectural condensation, suggesting a possibly separable step. Surprisingly, we show the small A-repeat fragment of XIST can alone silence endogenous genes; however, results indicate this requires high local RNA density for effective histone deacetylation. Results support a concept whereby XIST RNA acts directly to condense the chromosome territory, comprised largely of non-coding DNA, which facilitates a required step to initiate gene silencing by the A-repeat. Hence, compacted architecture is not a consequence of collective gene silencing, but an early step required for chromosome-wide gene silencing.

## INTRODUCTION

Transcriptional silencing of one X-chromosome in mammalian female cells is a pre- eminent model for the broader formation of facultative heterochromatin and epigenetic programming of early embryonic cells. Beginning in pluripotent cells, the X-linked human *XIST* or mouse *Xist* gene produces a long (14-17kb) noncoding RNA that coats one X-chromosome *in cis*, triggering a cascade of repressive histone modifications and chromosome-wide gene silencing; this process transforms the active chromosome territory to the condensed, heterochromatic Barr body (Creamer and Lawrence, 2017; Dixon-McDougall and Brown, 2016; Strehle and Guttman, 2020). Many studies have focused on how mouse Xist RNA recruits specific histone modifiations and how each contributes to the gene silencing process. Here our goal was to examine this for the *human* chromosome silencing process, but also to investigate key questions yet to be addressed or understood in either system. Notably, we investigate early steps of the process directly in the structural context in which XIST RNA functions: the nuclear architecture of a chromosome territory. The cytological-scale condensation of the chromosome has been commonly thought to reflect the collective effect of silencing individual genes across the chromosome. However, we earlier raised the “Barr body Paradox” to question why a large, cytological-scale dense body would be formed by local condensation of many silenced genes, given that active genes constitute only ∼10% of chromosomal DNA in a given cell-type (Hall and Lawrence, 2016). Furthermore, genes cluster on mitotic chromosomes and are non-randomly organized further within the geography of nuclear chromosome territories (reviewed in (Bickmore, 2013; Smith et al., 2020). Investigating chromosome silencing in direct cytological context provides a unique window into poorly understood but fundamental aspects of chromosome structure and epigenome regulation. Hence, here we investigate human XIST RNA’s role in initiating canonical gene silencing in relation to not only various histone modifications, but directly in relation to *in situ* chromosome architecture.

XIST expression and function begins in pluripotent cells of the inner cell mass and thus is best investigated in that context. However, human female ES or iPS lines have a precociously silenced X-chromosome or other anomalies (Diaz Perez et al., 2012; Hoffman et al., 2005; O’Neill et al., 2003; Tchieu et al., 2010), and even “naïve” female hESC lines do not reliably recapitulate full chromosome silencing (Sahakyan et al., 2017). Given lack of more tractable human systems, the field has relied heavily on mouse ES cells (or embryos), although there are known difference in XIST/Xist transcript structure and transcriptional regulation (Chow and Brown, 2003; Migeon, 2017). A chromosome silencing process that takes ∼4-5 days in differentiating female mESC has been described (Chaumeil et al., 2006). Since the silent state becomes irreversible and “XIST-independent” after several days of ES cell differentiation (Jiang et al., 2013; Pandya-Jones et al., 2020; Wutz et al., 2002), we compared the process in human iPS cells maintained as pluripotent or allowed to differentiate. Despite lists of chromatin modifications and a few validated XIST RNA binding proteins, it remains largely unknown what actually silences gene transcription and modifies architecture (Brockdorff et al., 2020). It is critical to identify the changes that most immediately follow expression and spread of XIST RNA. Therefore, much of this study focuses on events within 2-4 hours of XIST induction, as we sought to define inter-relationships between early XIST RNA distribution, specific protein modifications, chromosome territory condensation and nuclear position, and canonical gene silencing.

To enable investigation of the *human* chromosome silencing process in appropriate developmental cell context, this study capitalizes on a human iPSC system carrying a single inducible XIST cDNA (14kb) targeted into one trisomic chromosome 21. This system was previously shown to silence genes chromosome-wide and induce Xi epigenetic hallmarks on a chromosome 21 “Barr body” (Czerminski and Lawrence, 2020; Jiang *et al*., 2013). Silencing a trisomic chromosome circumvents creation of a functional monosomy or nullisomy that selects against XIST function, as occurs in several mouse Xist transgene studies. Our inducible human XIST cDNA lacks regulatory sequences that control XIST transcription, allowing focus on events downstream of synchronous XIST expression to examine RNA function.

The first part of this study focuses on full-length XIST RNA, with results that show the importance of understanding how initiation of canonical gene silencing relates to architectural changes to the chromosome. To gain further insights, we created an inducible XIST “minigene” carrying just the conserved XIST “A-repeat” domain (∼450bp), inserted in the same location as full-length XIST in the parental trisomic iPS cells. Numerous mouse studies have established that deletion of this tiny region from the much longer XIST transcript prevents gene silencing (Bousard et al., 2019; Colognori et al., 2019; Ha et al., 2018; Wutz *et al*., 2002). However, a distinct question is whether just this small A-repeat fragment can alone silence endogenous chromosomal genes, outside the context of 96% of the XIST transcript. One study examined this in a somatic cell line and concluded that silencing of endogenous genes required other XIST domains (Minks et al., 2013), however here we re-examine this question in an iPS cell context that more fully supports full-length XIST RNA function.

We investigate early distribution and function of XIST RNA in relation to changes induced at three levels: biochemical, structural and transcriptional. Hallmark histone modifications that become strongly enriched upon chromosome silencing include H2AK119ub and H3K27me3 (via PRC1 and PRC2, respectively), as well as H4K20me and macroH2A. Importantly we also examine a non-histone nuclear structural protein, CIZ1 (Ainscough et al., 2007), and subsequently consider histone H3K27 deacetylation in relation to the function of the A-repeat. Architectural parameters examined include XIST transcript distribution and territory coalescence, DNA condensation and Barr body formation, as well as chromosome relocation to the nuclear periphery. Allele-specific silencing of canonical genes is examined by RNA FISH, as is formation of the “CoT-1 RNA hole”, a region lacking heterogeneous nuclear RNA that overlaps the Barr body.

Analysis of multiple parameters on single inactivating chromosomes allows identification of precise temporal and spatial relationships between XIST RNA density distribution and other specific changes. Results support a new model whereby XIST RNA functions along two distinct tracks: one that acts more directly to modify architecture, in addition to the established track of recruiting histone modifiers to silence active genes. Among several new findings, we show that the deacetylase function of the small A-repeat domain is both necessary and sufficient to initiate local gene silencing, but this step is density-limited; other XIST domains are required to propagate over the chromosome and coalesce a dense XIST RNA territory on a compacted nuclear chromosome.

## RESULTS

To investigate the interrelationships between spread of XIST RNA and changes to overall architecture, histone modifications, and transcriptional silencing, we examined RNA, DNA and proteins on individual inactivating chromosomes in human iPS cells using molecular cytology. The importance of understanding the RNAs relationship to chromosome architecture is best appreciated by recognizing the initial size of the decondensed chromosome territory across which XIST transcripts must spread and the magnitude of overall condensation induced by XIST RNA. Although the Xi DNA territory, as usually observed in somatic cells, appears only modestly smaller than the Xi (visualized with a whole X-chromosome DNA library) (Fig 1A), the true scale of chromosome compaction enacted by XIST needs to be understood in the context of pluripotent cells in which XIST RNA expression/function begins, and which have much more decondensed chromatin than somatic cells. For example, in human H9 ES cells, which contain a precociously inactivated X-chromosome (Hall et al., 2008), there is a dramatic difference in size between a highly distended Xa-chromosome territory and the compacted Xi-territory (Fig1B). This point emphasizes the extent to which the initiation process requires not only that *XIST* repress transcription of genes, but this unique lncRNA must function across broad physical space to enact large-scale structural transformation. This point provides perspective for other observations below.

**FIGURE 1:**
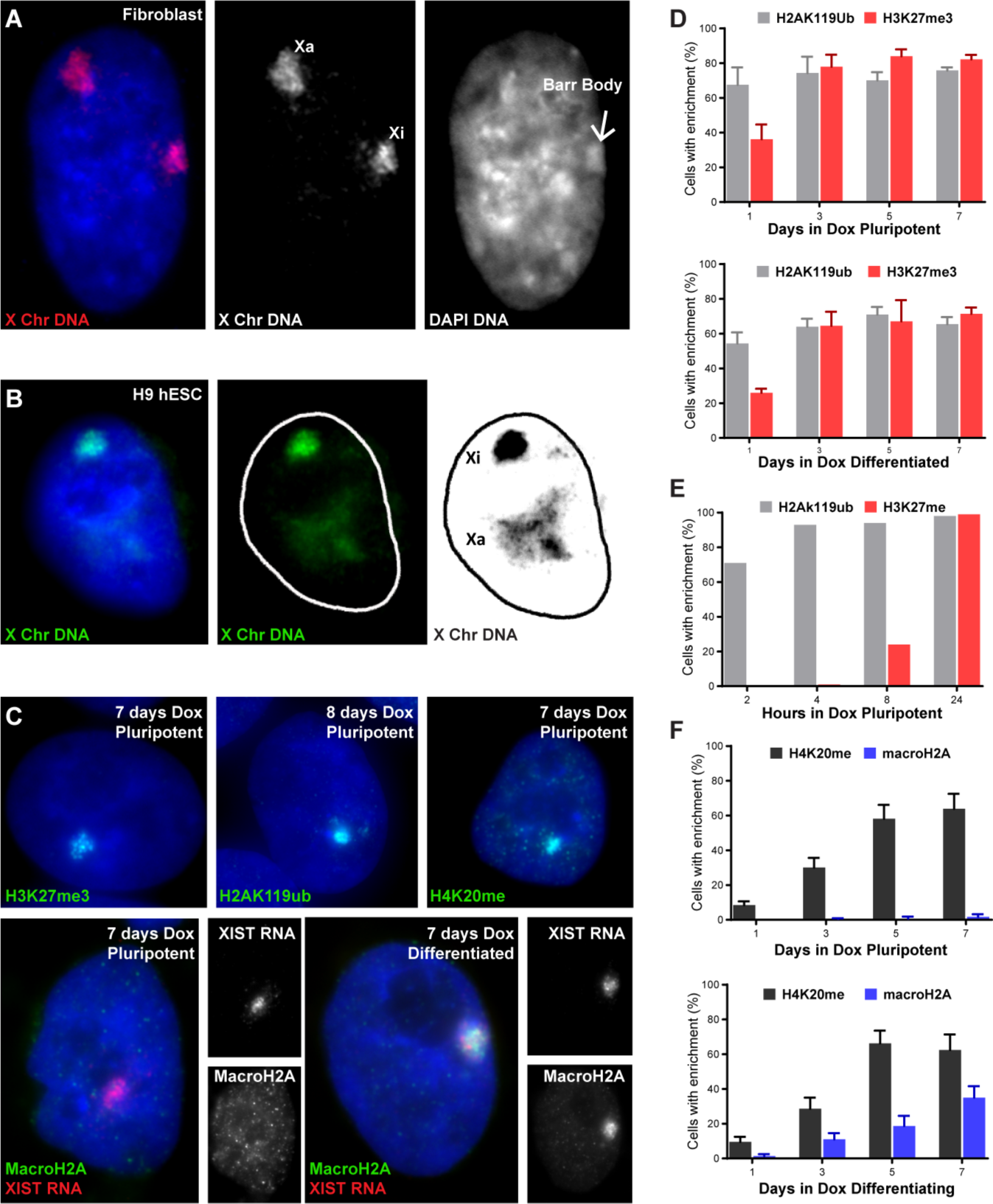
Human XIST RNA compacts a highly distended chromosome while heterochromatic hallmarks are sequentially accumulated. DAPI DNA is blue (all images). **A)** Active (Xa) and inactive (Xi) X-Chromosomes in somatic nucleus labeled with X-Chromosome paint. Red and Blue channels separated at right, with DAPI dense Barr body indicated (arrow). **B)** Same chromosome paint in pluripotent nucleus. Green channel with nuclear outline at right and inverted black & white far right. **C)** Immunofluorescence (IF) alone or with XIST RNA FISH for 4 classic heterochromatin hallmarks after 1 week of XIST expression in pluripotent or differentiated iPSCs. **D-E)** Enrichment of H4K27me3 & H2AK119ub by IF, after 1-7 days (D) or 2-24 hours (E) of XIST expression. **F)** Enrichment of H4K20me & macroH2A by IF, after 1-7 days of XIST expression.

### Human XIST RNA triggers H2AK119ub within two hours followed by H3K27me3, H4K20me, and macroH2A

To examine steps in the initiation of human chromosome silencing with high temporal resolution we used our XIST-transgenic trisomic iPSC system to synchronously induce XIST RNA. Previous transcriptomic analyses in pluripotent and differentiated cells showed comprehensive silencing of the ∼300 genes across chromosome 21 *in cis* (Czerminski and Lawrence, 2020; Jiang *et al*., 2013), and this is accompanied by compaction of the initially distended Chr 21 territory (Suppl Fig 1A). While those studies examined cells induced for a longer time (20 days), to study the initial steps of XIST function we began by examining the appearance of four canonical heterochromatin hallmarks after 1-7 days of dox-induction of XIST RNA. Immunofluorescence assays for H3K27me3, H2AK119ub, H4K20me and macroH2A each produce a bright signal against the darker nuclear background (Fig 1C), allowing sensitive visualization of these marks and XIST RNA on the same chromosome. Since XIST RNA expression begins in pluripotent cells just prior to differentiation, we initially compared the process in cells maintained as pluripotent or in those allowed to differentiate after dox induction, which would reveal if timing or presence of any of these modifications is differentiation-dependent.

As shown in Fig 1, enrichment for H4K20me and macroH2A appears days after H2AK119ub and H3K27me3 (Fig 1D&F). Interestingly, all marks appeared with similar kinetics in pluripotent versus differentiating cells except for macroH2A, which generally accumulated after the switch to differentiation conditions (see also: Suppl Fig 1B-G). Even in differentiating cultures, macroH2A lagged H4K20me by generally two days indicating macroH2A occurs later and is more differentiation dependent (Fig 1F); we note that some variability in the timing of macroH2A was seen and may reflect methods of iPSC culture and maintenance (see Methods & Suppl Fig 1H).

Both H2AK119ub and H3K27me3 accumulate on the inactivating chromosome in many cells by Day 1 and reached maximum by Day 3, independent of differentiation (Fig 1D). It is important to know which of these marks are recruited first by human XIST RNA, since earlier reports in mouse suggested Xist RNA recruits PRC2 first (for H3K27me3), followed by PRC1 (for H2AK119ub)(Zhao et al., 2008), reflecting their canonical relationship, while subsequent reports suggest initial deposition of H2AK119ub on Xi occurs before H3K27me3 (Almeida et al., 2017; Zylicz et al., 2019). We therefore examined cells just 2, 4, and 8 hours after adding doxycycline, and scored H2AK119ub and H3K27me3 enrichment in XIST expressing cells (on parallel slides in the same experiment). Results demonstrate a clear temporal difference, with H2AK119ub remarkably quick, and enriched in 71%, 93% and 98% of XIST RNA-positive cells at just 2, 4, and 8 hours, respectively (Fig1E). In contrast, in parallel samples only ∼24% of cells accumulated H3K27me3 by 8 hours, and similar results in multiple *in situ* experiments affirmed this order. We conclude that in human cells XIST RNA triggers strong H2AK119ub modification by PRC1 several hours *before* H3K27me3.

The appearance of H2AK119ub at the earliest time, just two hours after adding doxycycline, shows extremely close temporal connection with the initial onset of XIST RNA expression.

### Sparse XIST transcripts trigger H2AK119ub in broad territory whereas H3K27me3 is initially confined to smaller dense zone

We visualized XIST RNA spread across the inactivating chromosome territory relative to both the temporal appearance and spatial distribution of H2AK119ub and H3K27me3, at very early time points. XIST RNA first forms a very small intense transcription focus (Fig 2A), but sensitive RNA FISH analysis also consistently detects very low levels of XIST transcripts that spread much further within 2-4 hours, but localize within a discrete large nuclear territory (Fig 2B & F and Suppl Fig 2A). As explained under Methods, these low-level transcripts are visible through the microscope by eye, but may be missed if hybridization conditions (or digital imaging) are not optimal. By 8 hours many cells show this sparse punctate distribution of XIST transcripts in a larger region surrounding a smaller bright focal center of high-density RNA (encompassing the transcription focus) (Fig 2C). We will refer to these two regions of differing XIST RNA density as the “sparse-zone” and “dense-zone”. Importantly, we detect similar distribution of Xist-RNA in dense and sparse zones during the early stages of chromosome silencing in Xist-transgenic mouse ES cells (Fig 2E) and in very early mouse embryos during X-inactivation (Suppl Fig 2C). We note that the very low-level regional spread of human XIST RNA shown here is distinct from complete dispersal of XIST RNA throughout the entire nucleus, as illustrated when the RNA is released to randomly drift from the interphase chromosome by brief (4 hour) treatment with tautomycin (Hall et al., 2009) (Fig 2G).

**FIGURE 2.**
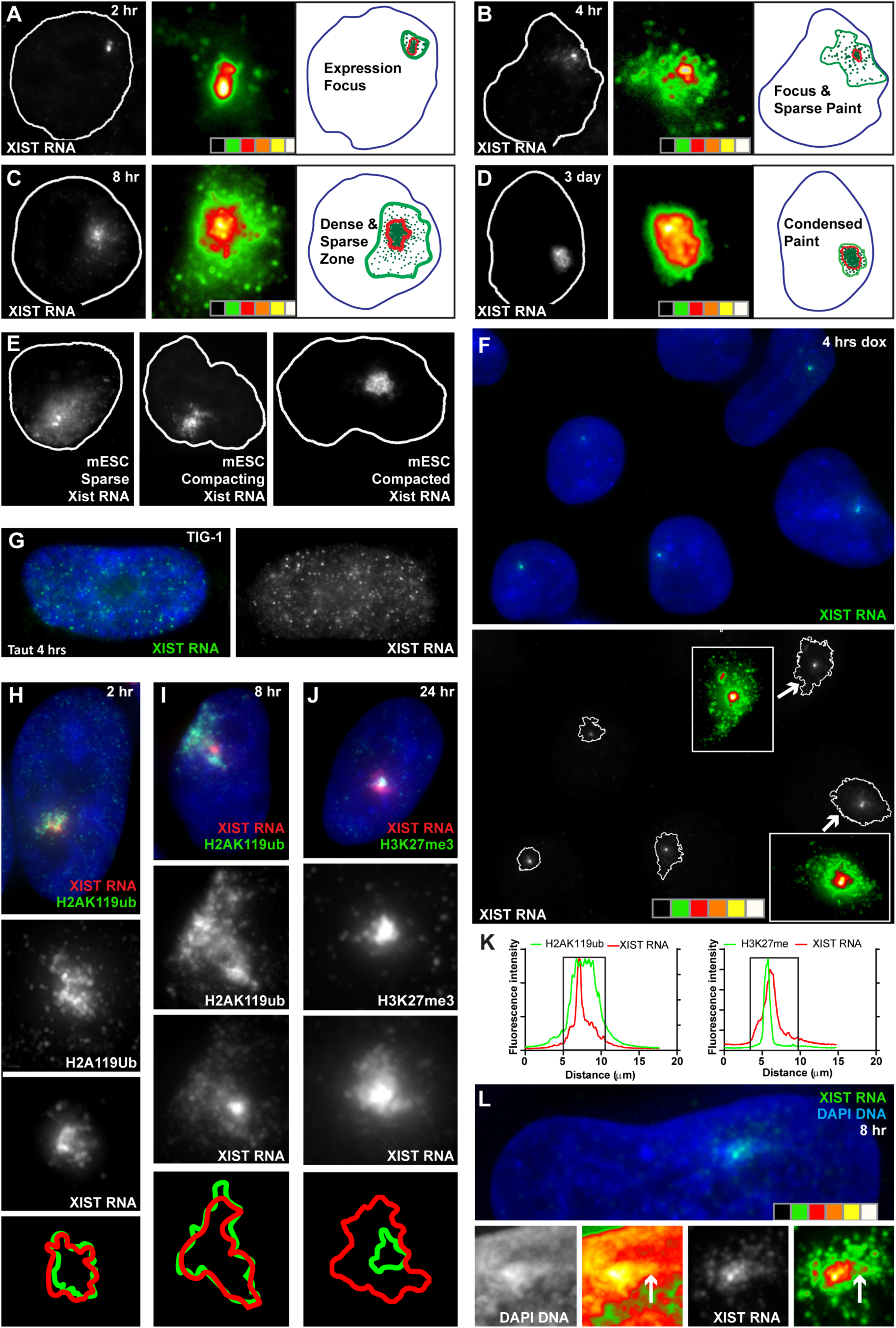
XIST RNA spreads broadly at low density within hours and alters chromatin differently when at high and low density. DAPI DNA is blue (F-L). **A-D)** XIST RNA (FISH) in nuclei over time-course of XIST expression. Black & white image shows RNA signal with outline of nucleus. Heatmap of XIST RNA signal intensity at center and illustration showing sparse (green) and dense (red) XIST RNA zones in nucleus (blue) at right. **E)** Xist RNA (FISH) territories over time-course in differentiating mouse ES cells containing an inducible Xist transgene integrated on Chr11. Black & white image best shows the early sparse RNA territory, with outline of nucleus. **F)** A field of cells after 4 hours XIST induction. Green channel is separated below with edge of XIST sparse-zone signal-threshold outlined in white. Select foci are shown as heatmap (inserts) to illustrate density changes. (Larger fields in supplement.) **G)** Tautomycin treated human Tig-1 fibroblasts release and fully disperse XIST RNA. Green channel is separated at right. **H-J)** IF of H2AK119ub (H-I) and H3K27me3 (J) with XIST RNA FISH. Red & green channels separated below, with edges of signal-threshold outlined at bottom. K) Linescans of representative nuclei showing IF labeling across the XIST RNA territory (boxed region) for H2AK119ub and H3K27me3 after 8 hours Dox induction. **L)** DAPI condensation in dense XIST RNA zone. Separated channels and representative intensity heatmaps of the XIST RNA region in close-up, below.

Since H2AK119ub and H3K27me3 enrichment both appear early, we examined their distribution relative to XIST RNA on individual chromosomes. The tight temporal connection between XIST RNA and H2AK119ub is further reflected in their relative distributions. Notably, H2AK119ub is elevated throughout the whole XIST RNA territory including the large sparse-zone (Fig 2H-I & K and Suppl Fig 2D). Even at just two hours when we can see a very low level of XIST transcripts in the sparse-zone, this is coincident with clear, often bright, enrichment for H2AK119ub. In contrast, H3K27me3 is incorporated not only later (shown above) but is much more restricted to the smaller dense XIST RNA zone (Fig 2J & K). Thus, H2AK119ub staining mirrors XIST RNA distribution largely independent of density, while H3K27me3 enrichment is limited to the dense-zone. If RNA hybridization is omitted (to rule out any impact of hybridization procedures), H2AK119ub again clearly and consistently marks a region larger than that of H3K27me3 (Suppl Fig 2E).

Importantly, the low levels of sparsely distributed XIST RNA shown here are neither noise nor inconsequential “drift”, but transcripts functionally interacting with chromatin, as evidenced by appearance of H2AK119ub. The striking speed of this is consistent with other evidence that PRC1 complexes are already present (Chu et al., 2015; Nesterova et al., 2019; Zylicz *et al*., 2019). Most importantly, these findings point to differences in XIST transcript *density* as a significant parameter, which influences its functional effects, indicating distinct histone modifications differ in the requirements for transcript density.

Between ∼1-3 days following XIST induction the dense RNA zone expands and encompasses the progressively contracting sparse-zone. Ultimately, the more compact and uniformly dense XIST RNA territory is formed (e.g. Fig 2D & Suppl Fig 2B), as typical of XIST RNA on the compacted chromosome of somatic cells. At early time points, the small dense RNA zone (which overlaps H3K27me3) is often coincident with a focal increase in DNA condensation (Fig 2L), indicating an early stage in nucleation of the Barr body, as further examined below. Less frequently slightly increased DAPI-DNA density is evident across the larger XIST RNA sparse-zone (Fig 2L arrow) but only discernible using optical sectioning and deconvolution for high-resolution maximum image projections. More readily and consistently observed was that XIST RNA rapidly spread very sparsely across a decondensed territory and subsequently transcripts cluster and merge to build a dense RNA territory coincident with compaction of the chromosome.

### XIST RNA acts early to modify architecture before silencing most protein-coding genes

The mature Barr body of somatic cells has been shown to comprise a dense core of repeat-rich silent DNA, readily delineated by a void in Cot-1 RNA signal *in situ* (e.g. Suppl Fig 3A) (Clemson et al., 2006; Hall et al., 2002). Hence, we examined CoT-1 RNA as a hallmark for architecture, and to compare formation of this dense silent domain to temporal silencing of canonical genes. The Barr body was long thought to comprise the whole Xi (condensed due to gene silencing), however, it was discovered that all of 14 genes examined positioned just outside the DNA-dense Barr body, in the periphery of the XIST RNA territory (Clemson *et al*., 2006); similar peripheral localization can be seen on the Xi in mESCs (Chaumeil *et al*., 2006). Moreover, on active chromosomes genes mostly distribute in the peripheral zone (Bickmore, 2013; Bickmore and Teague, 2002; Clemson *et al*., 2006; Kurz et al., 1996; Mahy et al., 2002; Scheuermann et al., 2004), often with inter-chromosomal domains enriched for splicing factors (Xing et al., 1993; Shopland et al., 2003, Chen and Belmont, 2019; Smith et al., 2020). Hence, this spatial separation of genes within the chromosome territory suggest that gene silencing may also be temporally separable from architectural formation of the Barr body, marked by a void in CoT-1 RNA (Hall and Lawrence, 2010).

RNA FISH was used to examine the temporal and spatial relationships of specific gene silencing to formation of the Cot-1 RNA depleted Barr body. A region depleted of CoT-1 RNA was generally seen with XIST RNA by day 1, therefore we examined shorter time-points (Fig 3A-B). A region with modest depletion of CoT-1 RNA could be discerned in some cells at two hours, which became more evident at 4 and 8 hours (Fig 3B & Suppl Fig 3B-C). The CoT-1 RNA depleted region was often clearest at the small dense-zone of brightest XIST RNA, with less apparent depletion over the sparse-zone; this pattern is reflected in the “V” shape of the linescan (Fig 3B). By 24 hours a more clearly defined larger region of low CoT-1 RNA is seen, which becomes darker and eventually encompasses the XIST RNA territory by Day 3 (Suppl Fig 3D), although in some cells the “CoT-1 hole” appears more fully formed at later time points.

**FIGURE 3.**
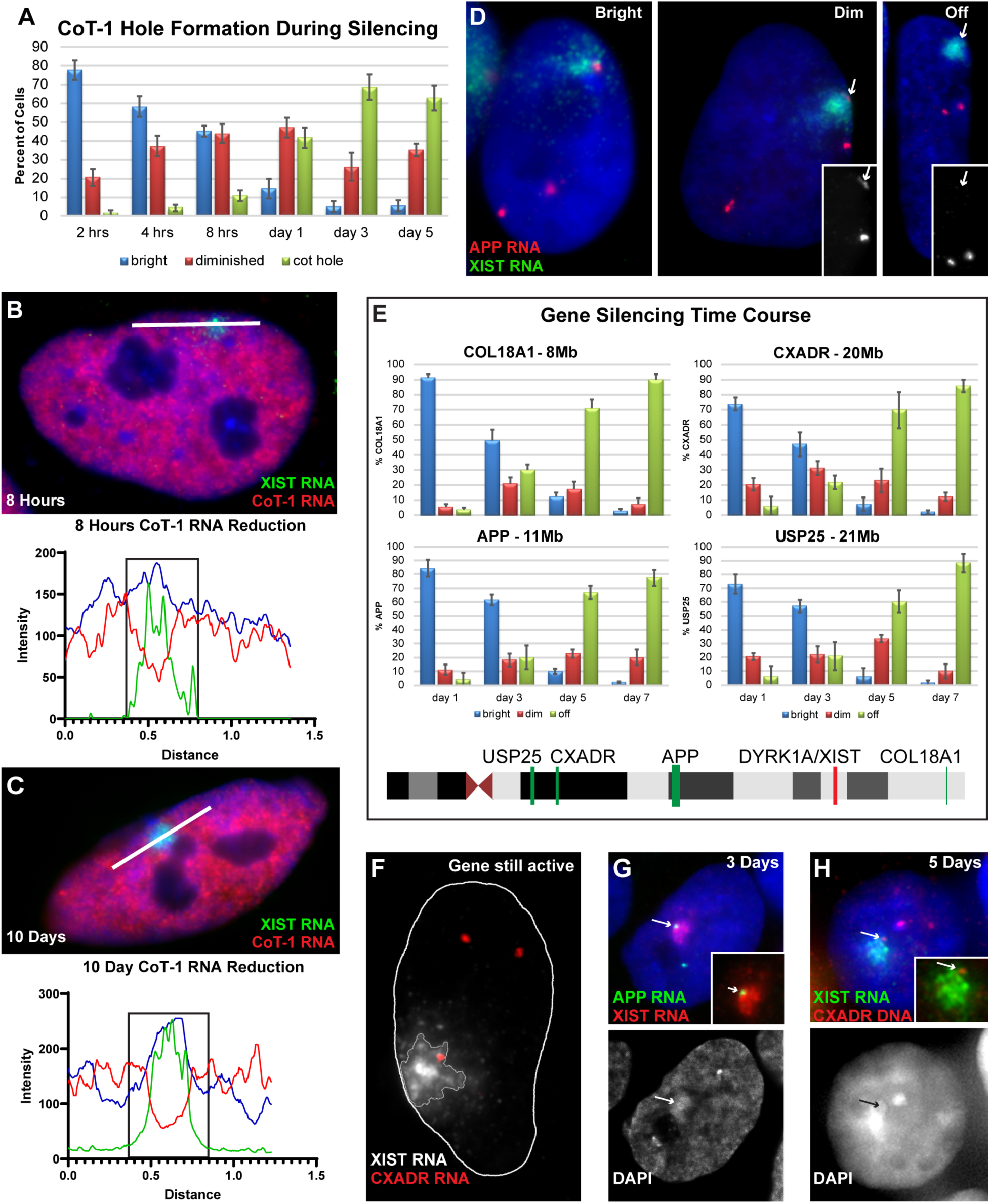
Formation of Barr body architecture occurs days before most gene silencing. DAPI DNA is blue (B-D & G). **A)** CoT-1 RNA hole formation (RNA FISH) over time-course of XIST expression in pluripotent iPSCs. **B-C)** CoT-1 and XIST RNA FISH with representative linescans below. Edges of XIST RNA signal indicated by black box. **D)** XIST and APP RNA FISH in differentiating iPSCs. Inserts: single channel close-up of APP gene signals (arrows). **E)** Quantification of gene silencing (loss of transcription focus) for four genes over time-course of XIST expression in pluripotent iPSCs. Ideogram of gene location on Chr21 below. **F-H)** Illustrate early and later gene positioning and silencing relative to XIST RNA territory and Barr body. **F)** CXADR and XIST RNA FISH. Outline of nucleus in white and threshold of XIST RNA signal also outlined. **G)** XIST and APP RNA FISH. DAPI channel separated below, with APP transcription focus relative to XIST RNA/Barr body indicated (arrows). Inset: APP and XIST RNA with blue channel removed for clarity. **H)** XIST RNA and CXADR DNA FISH. DAPI channel separated below, with location of CXADR gene relative to XIST RNA/Barr body indicated (arrows). Inset: CXADR gene (DNA) and XIST RNA with blue channel removed for clarity.

As we previously showed (Clemson *et al*., 2006; Xing et al., 1993), RNA density for any gene is highest at the site of transcription, hence RNA FISH analysis of transcription foci provides a direct read-out of allele-specific gene silencing on the XIST RNA coated chromosome. We identified genomic probes that detect with high efficiency pre-mRNA foci for four genes which map widely across the chromosome (8-21 MB from *XIST*). We quantified silencing at days 1, 3, 5, and 7, with CoT-1 RNA examined in parallel. While a CoT-1 RNA depleted domain was apparent in most cells by Day 1 (e.g. Fig 3A), at this time point none of the four genes showed any loss (or reduced intensity) of transcription from the XIST associated allele compared to the other two alleles in the same cell (Fig 3D-E). Importantly, transcription foci for all four genes continue to be synthesized in these rapidly dividing cells, with no significant silencing observed until Day 3 of XIST expression, and this was not maximal until Day 7, in either pluripotent or differentiated cells.

Figure 3F further illustrates that transcription foci for these genes are expressed from peripheral regions, in the larger sparse-zone of the XIST RNA territory, away from the denser RNA zone that forms near the XIST transcription site. In keeping with the organization shown for numerous Xi genes in human fibroblasts (Clemson *et al*., 2006) and differentiating mouse cells (Chaumeil *et al*., 2006), by Day 5-7 silenced genes have come “inward”, but still distribute primarily in the peripheral rim of the condensed chromosome and coalesced RNA territory (Fig 3G-H). In sum, it is remarkable that the large DAPI dense domain lacking CoT-1 RNA (Barr body) is essentially formed about two days before long-range gene silencing occurs.

### XIST rapidly impacts CIZ1 architectural protein and silences genes before chromosome movement to the peripheral lamina

Most studies of XIST RNA function focus on the RNA recruiting histone modifiers to trigger a cascade of histone modifications, which are known to impact local chromatin structure at the nucleosomal level, and thus gene transcription. Larger-scale chromosome condensation, a hallmark of the chromosome silencing process, is commonly thought to reflect additive effects of chromosome-wide histone modifications and gene silencing. However, the above findings indicate that XIST RNA acts quickly to modify cytological-scale architecture, well before most gene silencing and multiple hallmark histone modifications. Hence, this suggests a fundamentally distinct alternative model for XIST RNA function which includes more direct impact on elements of larger-scale architecture. These alternative models (outlined in Figure 7) will be further considered in the Discussion, but motivated our next analysis of non-histone elements of chromosome structure identified as part of a nuclear matrix or scaffold.

In earlier work demonstrating that XIST RNA paints the Xi DNA territory, we showed that XIST RNA remains bound with the classically defined non-chromatin nuclear matrix (Clemson et al., 1996), and we recently affirmed and extended this finding using a more selective fractionation procedure (Creamer et al., 2021). Two matrix proteins, SAF-A (Helbig and Fackelmayer, 2003) and CIZ1 (Ridings-Figueroa et al., 2017; Sunwoo et al., 2017) have been shown enriched on Xi and thought to function as chromosomal tethers for XIST RNA so that the RNA can act *in cis* to trigger histone modifications. CIZ1, like SAF-A, has both RNA and DNA binding domains, and is thought to be recruited by XIST RNA in order to anchor RNA to the chromosome (Hasegawa et al., 2010; Kolpa et al., 2016; Ridings-Figueroa *et al*., 2017; Sunwoo *et al*., 2017). However, we consider here that their roles may go beyond anchoring RNA to the chromosome.

As shown in Fig 4A, immunofluorescence for SAF-A shows broad chromatin distribution in pluripotent cells (before any XIST expression), consistent with its ubiquitous expression in various cell-types (Helbig and Fackelmayer, 2003; Kolpa et al., 2016). In contrast to SAF-A, CIZ1 staining is essentially negative in iPSCs prior to XIST induction, with at most a few tiny puncta visible against a dark background (Fig 4B). However, in cells expressing XIST RNA, a very bright territory of CIZ1 overlaps the XIST RNA territory in an otherwise empty nucleoplasm (Fig 4B). Because robust CIZ1 signal was seen with XIST RNA at Day 1, we examined hourly time points and found many cells formed the bright CIZ1 territory within just two hours of adding doxycycline (Fig 4B-C).

**FIGURE 4.**
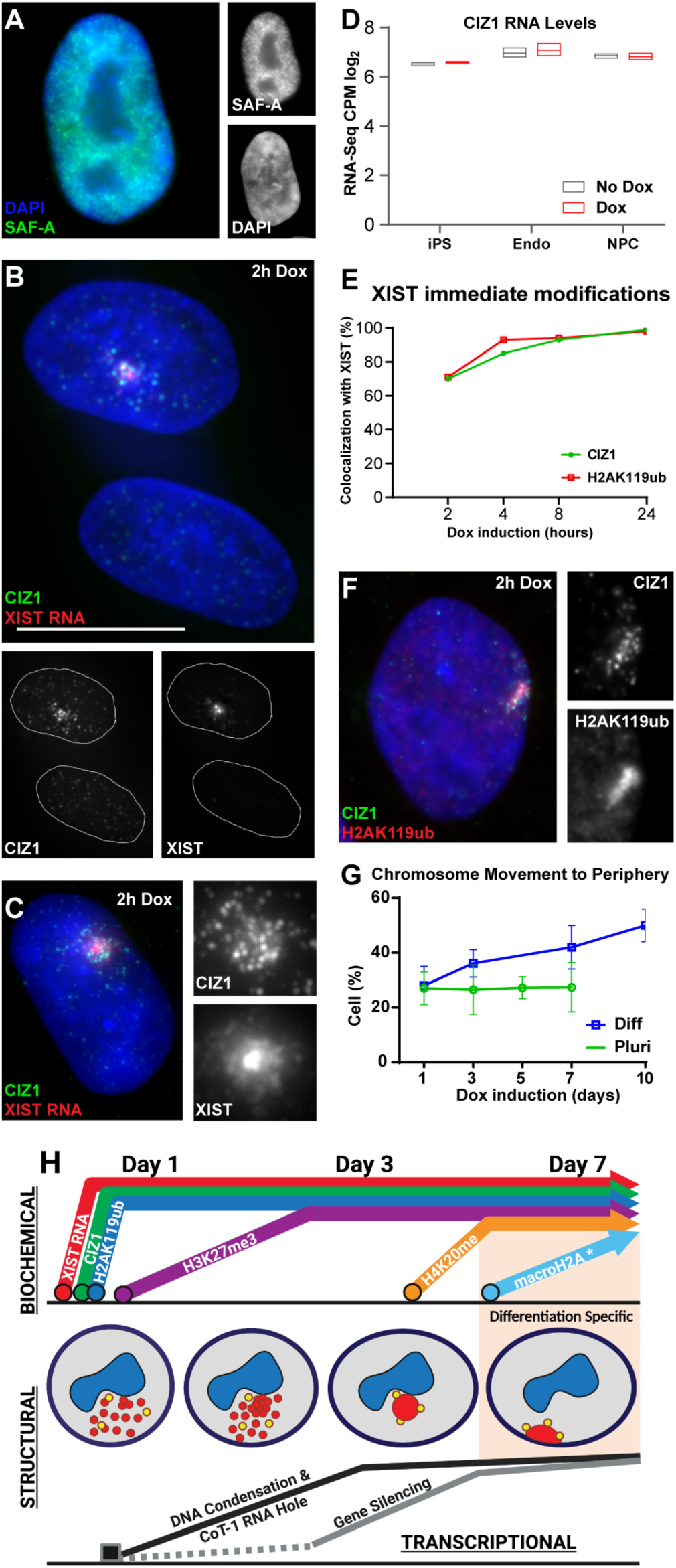
XIST RNA rapidly impacts CIZ-1 scaffold factor whereas chromosome peripheral movement is late and requires differentiation. DAPI DNA is blue (A-C & F). **A)** SAF-A IF in pluripotent iPSC nuclei. Separated channels at right. **B)** CIZ1 (IF) and XIST RNA (FISH) in cells expressing XIST for two hours compared to neighboring non-expressing cells. Separated channels below with nuclei outlined in white. **C)** CIZ1 (IF) and XIST RNA (FISH). Close-up of XIST RNA region with separated channels at right. **D)** CIZ1 mRNA levels in pluripotent and differentiated iPSCs (Endothelial & Neural progenitor cells). **E)** Scoring nuclei with XIST RNA (FISH) and CIZ1 or H2AK119ub enrichment (IF) hours after XIST induction. **F)** Simultaneous CIZ1 and H2AK119ub IF. Close-up of XIST RNA region with separated channels at right. **G)** Percentage of pluripotent and differentiated cells with transgenic Chr21 located at nuclear periphery. **H)** Schematic summary of results showing temporal order of numerous changes triggered by human XIST RNA on inactivating chromosome within nucleus (dark blue, nucleolus). Summary integrates analysis of parallel samples analyzed for changes at the biochemical, structural and transcriptional levels. Within 2-4 hours XIST RNA (red) spreads across large sparse zone, which coalesces to dense XIST RNA territory concomitant with condensation of highly distended DNA territory of pluripotent cells. Formation of a compartment lacking Cot-1 RNA marks condensation of repeat-rich DNA to form the Barr body. Timing of silencing is shown for the four genes examined (yellow), which occurs as genes initially distant from dense XIST RNA move inward to the border of a coalesced RNA territory, an organization seen previously for 14 genes on mature Xi (see text). The order and timing of steps was similar in pluripotent or differentiating cells, except movement of silent human chromosome 21 (or Xi in hESCs) was only seen after differentiation. MacroH2A enrichment also was typically seen after differentiation, with some exception (see text).

At all early time points CIZ1 is strongly detected in both the sparse- and dense-zones of XIST RNA, closely mirroring RNA distribution. Given the lack of CIZ1 staining in pluripotent cells (nuclei or cytoplasm) before induction, it was surprising that such a large, robust accumulation of CIZ1 appears so quickly, with no change in the negligible nucleoplasmic fluorescence. This very short-time frame is difficult to reconcile with XIST RNA inducing CIZ1 expression and subsequently recruiting newly synthesized protein. In support of this, we examined RNAseq data from iPSCs and endothelial cells generated for another study (Moon and Lawrence, submitted) and found that CIZ1 mRNA is clearly expressed in iPSCs irrespective of XIST induction, and only modestly higher post-differentiation (Fig 4D). Rather than XIST RNA recruiting CIZ1 to the chromosome, these results suggest that CIZ1 is already there but the epitope detected by a monoclonal antibody is masked in pluripotent cells, *except* when interacting with XIST RNA. Indeed, earlier studies focused on CIZ1’s role in DNA replication (Coverley et al., 2005) showed that CIZ1 is present broadly in nuclei but only detectable by IF (with two antibodies) after chromatin removal in a matrix protocol; hence it was concluded that the CIZ1 epitope is masked by interaction with DNA (Ridings-Figueroa *et al*., 2017; Swarts et al., 2018). Interestingly, CIZ1 is known to also bind DNA (Warder and Keherly, 2003) and the monoclonal antibody we used targets the zinc finger region.

In fact, it is known that XIST RNA interaction with SAF-A masks detection of concentrated SAF-A on the Xi, however the SAF-A epitope is revealed in a chromatin-depleted matrix prep or by antigen retrieval (or GFP-tagged SAF-A) (Helbig and Fackelmayer, 2003; Kolpa *et al*., 2016). Hence, collective evidence favors that interaction of CIZ1 with XIST RNA also “unmasks” an epitope to reveal CIZ1 already present in iPSCs. As considered in the Discussion, XIST RNA does not itself bind and directly modify histones, but the RNA does bind CIZ1 and SAF-A (Helbig and Fackelmayer, 2003; Ridings-Figueroa *et al*., 2017), and thus may directly impact conformation and/or chromatin interactions of these architectural proteins. This would be a fundamentally distinct role from acting as a tether to localize histone modifications (see Discussion).

Both CIZ1 and H2AK119ub rapidly appear with induction of XIST RNA in 70% of cells (and ∼100% by 24 hours) (Fig 4E-F). Using simultaneous staining to discern any potential temporal order to their appearance, CIZ1 was occasionally seen without H2AK119ub, although results were inconclusive as to whether the inverse can also occur (Suppl Fig 4A & legend). However, it is clear that changes to both CIZ1 and H2AK119ub are very rapid, essentially concurrent and immediate, and both are broadly induced in the sparse (and dense) zones of XIST RNA.

These results show that one of the first effects of XIST RNA is to impact the CIZ1 nuclear matrix protein. The lamins are also architectural proteins of the nuclear matrix, and the Xi is known to preferentially associate with the lamina at the nuclear periphery, as seen in ∼80% of human fibroblasts. Chromosome repositioning to the lamina may be mediated by XIST RNA’s interaction with the lamin-B receptor (LBR)(Chen et al., 2016). This study also reported that movement to the peripheral lamina was required for gene silencing, however we find in human pluripotent cells Chr21 genes are silenced without chromosome relocation to the nuclear periphery (Fig 4G & Fig 1C). The silenced chromosome does relocate to the nuclear periphery in many cells, but only upon differentiation (Fig 4G). To address the possibility that an autosome (carrying rDNA genes) might behave differently, we also examined several pluripotent female human ES cell lines bearing a precociously inactivated X-chromosome (Hall *et al*., 2008), and again, only upon differentiation did the Xi become more peripheral (Suppl Fig 4B-C). Thus, chromosome interaction with lamina architecture occurs after differentiation, and requires one or more factors expressed later in differentiated cells, such as lamin A/C(Butler et al., 2009), or possibly SMCHD1(Wang et al., 2018). Our results show early events and gene silencing occurred similarly with or without differentiation. However, gene silencing in this system only becomes irreversible upon XIST RNA removal after differentiation (Jiang et al., 2013), therefore peripheral lamina association may be important for the irreversible heterochromatic state.

Figure 4H summarizes our findings regarding biochemical, architectural and transcriptional changes triggered by full-length human XIST RNA during initiation of *human* chromosome silencing. Our collective findings all point to a larger theme: that within two hours XIST RNA spreads widely at low levels to immediately impact certain histone *and non-histone* chromosomal proteins and begins remodeling overall architecture days before most transcriptional silencing of genes.

### RNA from just the small XIST A-repeat can silence transcription of local endogenous genes

To better understand the relationship between architecture and gene silencing, we investigated a domain of XIST, the A-repeat. Numerous studies have affirmed in mouse that an Xist mutant deleted for just this small (450 bp) A-repeat domain can no longer transcriptionally silence genes, even though the RNA still spreads widely across the chromosome (Brockdorff, 2018; Colognori et al., 2020; Engreitz et al., 2013; Ha *et al*., 2018; Wutz *et al*., 2002). Hence it is well established that the A-repeat is required for silencing, but here we investigate the reciprocal question: whether the tiny (450 bp) A-repeat might itself be *sufficient* to transcriptionally repress endogenous loci *in cis*. One previous study examined this question in human HT1080 fibrosarcoma cells using qRT-PCR and found A-repeat RNA could partially repress the GFP reporter integrated on the same plasmid (7 kb separation), but, importantly, could not significantly repress even immediately proximal endogenous loci (100kb-3Mb away) (Minks *et al*., 2013). Hence, it was concluded that sequences within the missing 96% of full-length XIST RNA are required to support the A-repeat function in gene silencing.

However, since XIST RNA mediated silencing is substantially compromised in HT1080 cells (Hall *et al*., 2002; Minks *et al*., 2013), we investigated this question further in human pluripotent cells, where XIST RNA function is optimal. As shown in Fig 5A, we employed the same inducible promoter, insertion site, editing methodology (ZFNs) and iPS cells as was used for full-length (14kb) *XIST* (*flXIST*) (Jiang *et al*., 2013) to engineer cells for inducible expression of the 450bp A-repeat “nanogene” (lacking 96% of 14kb flXIST transgene). A red fluorescent protein (RFP) gene under a constitutive promoter (EF1α) was included downstream of the A-repeat, and correct targeted insertion (into Chr21 DYRK1A intron) was confirmed by two-color RNA FISH in uninduced cells (Fig 5B).

**FIGURE 5.**
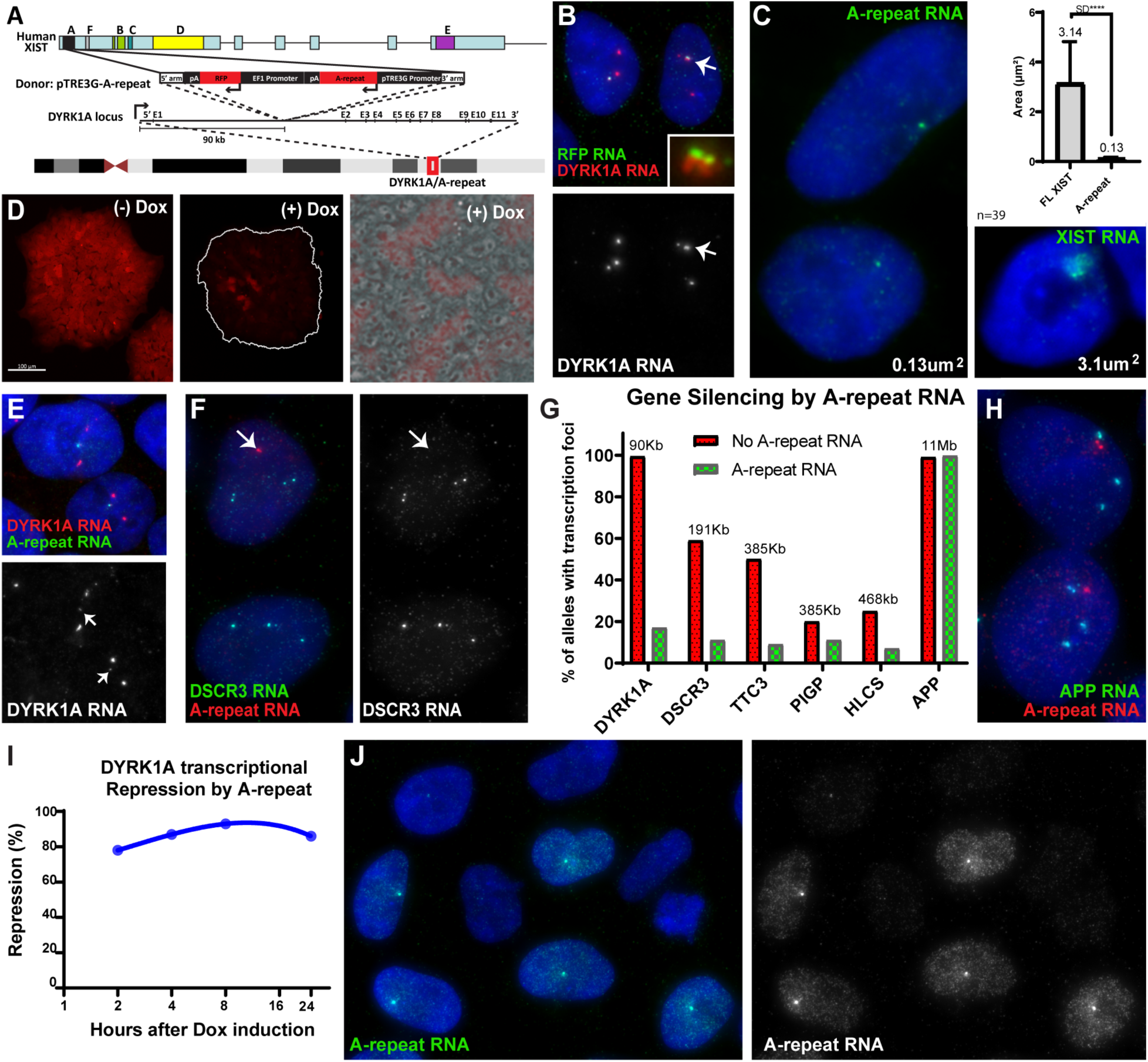
A-repeat RNA does not spread on chromosome but silences nearby endogenous genes as well as adjacent reporter. DAPI DNA is blue (all images). **A)** A-repeat transgene map and insertion site on Chr21. **B)** RFP and DYRK1A RNA FISH in uninduced cells, with co-localization of RFP and DYRK1A RNA at transgene target site (insert). Red channel separated below, with no reduction in linked DYRK1A TF (arrow). **C)** RNA FISH for indicated probes, with average RNA territory size indicated in image and in graph. **D)** RFP in transgenic iPS cell cultures with and without dox induction of A-repeat, showing loss of RFP with dox (middle, colony outlined in white). Close-up of representative Dox (+) colony (far right) indicates not all cells induce A-repeat. **E-F)** RNA FISH of indicated probes. Separated channels for Chr21-linked gene RNA below and at right. Locus with A-repeat transgene indicated (arrow). **G)** Quantification was performed from z-stacks of RNA FISH images. Frequency of un-linked alleles (red) versus those linked to A-repeat RNA (green). “Trace” signals for DYRK1A scored as silenced, due to slight read-through from transgene (Suppl Fig 5 for more details). **H)** APP and A-repeat RNA FISH. **I)** Quantification across time-course of repressed DYRK1A transcription focus associated with A-repeat. **J)** A-repeat RNA FISH showing substantial dispersed nucleoplasmic RNA in A-repeat expressing cells, adjacent to non-expressing cells.

Since it has not been examined previously, the distribution of A-repeat RNA was of interest. The A-repeat produced a much smaller but intense focal RNA accumulation, after dox induction, in clear contrast to the large flXIST RNA territory (Fig 5C).

Microfluorimetric measurements showed A-repeat RNA foci occupy an area ∼4-5% of the flXIST RNA territory, but the bright focal signal indicates high density of this small sequence at that site. Apart from this small focal accumulation, A-repeat RNA did not spread and localize substantially on the chromosome territory. Induction of A-repeat RNA was able to silence the RFP reporter gene integrated with the same plasmid under a separate promoter (1.7kb away), as iPS cell colonies began losing red fluorescence (Fig 5D), supporting results of (Minks *et al*., 2013). In experiments below, we used a subset of cells that do not induce A-repeat RNA (due to stochastic silencing of the tet-activator, see Methods) (Fig 5D & J) as a negative internal control for direct comparison of cells with and without A-repeat RNA (see below).

RFP and A-repeat transgenes are directly adjacent on the same inserted plasmid, but a distinct question is whether the 450 bp A-repeat transcript, expressed from an intron of a large gene (DYRK1A), can impact expression of that gene’s endogenous promoter (90kb away), and potentially other nearby endogenous loci (map in Suppl Fig 5A). To evaluate A-repeat effects on transcription, we used RNA/RNA FISH with gene-specific genomic probes to directly visualize transcription foci, which allows allele-specific analysis in single cells. Uninduced cultures show three clear DYRK1A RNA foci in essentially all cells, given high detection efficiency for this probe/RNA, (affirming the transgene in the intron does not affect DYRK1A transcription) (Fig 5B). After dox-induction for eight days, transcription foci from the DYRK1A allele *in cis* with the A-repeat were essentially silenced (83% of cells) (Fig 5E & G and Suppl Fig 5B), whereas normal bright transcription foci were maintained at the other two loci. Generally, transcription foci at A-repeat expressing loci were entirely absent or a barely visible trace (which other observations indicate is read-through from XIST into the DYRK1a intron, see Methods). For an extremely close tandem reporter gene (RFP in this study or GFP in (Minks *et al*., 2013)) it is harder to rule out that A-repeat effects are via steric hindrance, but the DYRK1A promoter is 90kb away. Furthermore, uninduced cells expressing bright RFP transcription foci had no repressive effect on the nearby DYRK1A promoter (only 8% showed smaller DYRK1A RNA foci, consistent with random variation) (Fig 5B). These results provided the first evidence that just the small A-repeat sequence itself retains gene silencing function and thus can repress a nearby endogenous locus.

Therefore, we next examined two other nearby genes that map significantly further from the integration site, DSCR3 (191kb away) and TTC3 (385kb away) (Suppl Fig 5A), which prior microarray results indicated are expressed in these iPSCs (Jiang *et al*., 2013), and for which we could generate appropriate genomic probes. Since the strength of transcription focus signals will vary for a given gene based on size, intron content, and expression level, three transcription foci are not as consistently detected in each cell as they are for DYRK1A. Hence, we first quantified transcription focus detection efficiency, using simultaneous detection of DYRK1A RNA foci for comparison (which also confirms the specific locus) (Suppl Fig5C-D). Detection frequencies of transcription foci for DSCR3 and TTC3 at each allele was 59% and 50%, respectively (Fig5F-G). While not our focus here, we note that the detection of transcription foci at two or all three alleles in many cells argues against single-cell seq analysis interpreted to show that most genes express from just one allele (see Suppl Fig5H and legend). Analysis of parallel dox-induced samples clearly showed silencing of the A-repeat associated allele (Fig5G and Suppl Fig5B-D), with the frequency of transcription foci dropping by 82% for DSCR3 and 83% for TTC3. This demonstrates that A-repeat RNA effectively repressed transcription of genes a few hundred kb away.

Given these surprising results, we worked to evaluate two other nearby expressed genes, PIG1 (385kb away) and HLCS (468kb away), for which transcription foci were detected at lower but significant frequencies (Suppl Fig5E-F). Nonetheless, frequency of transcription foci at the A-repeat allele dropped to 11% and 7% for PIG1 and HLCS, respectively, representing a drop of ∼46-74% (Fig5G). While the efficiency of A-repeat RNA silencing appears to decline across a ∼400 kb interval, silencing still occurred in many cells for loci as much as 438 kb away. We then also examined the more distal APP gene (11 Mb away). For this large gene, three transcription foci were always detected even after inducing the A-repeat, with no reduction in size or intensity of RNA from the allele nearest the A-repeat RNA foci (Fig 5G-H and Suppl Fig5G) (7% appeared smaller, consistent with modest stochastic variation). We conclude that, in the appropriate developmental cell context, just this small A-repeat fragment alone can silence transcription of *endogenous* genes. Importantly, this is limited to the “local chromosomal neighborhood” shown here for a region 400-450kb from the transcription site, with no effect on the APP locus several mega-bases away. Surprisingly, this 450 bp fragment retains this functionality outside the context of 96% of the XIST transcript.

Since the small A-repeat transcripts do not spread along the chromosome, a focal concentration may form rapidly. To test this and determine how long it takes the A-repeat RNA foci to silence local gene transcription, we induced cells for just two hours and examined levels of A-repeat and DYRK1A RNA. Within two hours of adding doxycycline, dense foci of A-repeat transcripts had formed in many cells, and in parallel had quickly repressed DYRK1A transcription foci from that allele (Fig 5I). Thus, this dense focal concentration of A-repeat RNA can very quickly silence nearby gene transcription.

In addition to the concentrated A-repeat RNA focus, many cells had A-repeat transcripts dispersed uniformly throughout the nucleoplasm (Fig5J), but these were not found in the cytoplasm, as was RFP mRNA, suggesting A-repeat transcripts are not exported (Suppl Fig 5I-J). Full-length XIST RNA is highly stable, with a half-life of about five hours (Clemson et al., 1998; Clemson *et al*., 1996); in contrast we found the A-repeat RNA focus dissipates after 30 minutes of transcriptional inhibition, and nucleoplasmic transcripts become undetectable after about an hour (Suppl Fig 5K). Hence, the A-repeat transcript accumulates locally to silence nearby genes but is released from chromatin to disperse. And although it is much less stable than flXIST, it is not immediately degraded (or transported) and thus can populate the nucleoplasm, as will be further considered below.

### Effective deacetylation to initiate gene silencing requires high density of A-repeat/XIST transcripts

Results above show that flXIST RNA spreads rapidly across the chromosome and that A-repeat RNA itself can silence genes, and can do so rapidly. Since flXIST contains the A-repeat and spreads across the chromosome territory within hours, why didn’t flXIST RNA induce long-range gene silencing more quickly? The widely distributed flXIST RNA is clearly sufficient to trigger robust H2AK119ub and CIZ1 staining within just two hours, yet it took days to silence several randomly selected genes, which occurs after coalescence of the XIST territory.

To gain insight into this we further considered how the A-repeat functions, since this sequence is required for the gene silencing process. This has been previously studied by deletion of the A-repeat, and two main functions have been implicated: histone deacetylation and chromosome organization with the nuclear lamina. Evidence indicates the A-repeat is required to recruit HDACs for H3K27 deacetylation (via SPEN) which is important in the chromosome silencing process (Brockdorff *et al*., 2020; Chu *et al*., 2015; McHugh et al., 2015; Nesterova *et al*., 2019; Zylicz *et al*., 2019). In addition, the A-repeat region binds the lamin B receptor (LBR) (McHugh *et al*., 2015) and the consequent tethering of the chromosome to the peripheral lamina was reportedly required for gene silencing (Chen *et al*., 2016). However, as shown above, our results with flXIST RNA do not support a requirement of peripheral localization for initial gene silencing. Likewise, A-repeat RNA foci do not localize to the nuclear periphery (in pluripotent or differentiated cells, Suppl Fig 6A), yet are still able to silence genes locally (e.g. Fig 5G).

We next investigated whether this 450 bp fragment of XIST acts via deacetylation to stop transcription. A 4-hour TSA treatment was sufficient to inhibit histone deacetylation and increase H3K27ac across the nucleoplasm (Suppl Fig6B-C), while short enough to avoid toxic effects. Gene silencing by either the A-repeat or flXIST RNA dropped markedly when histone deacetylation is blocked concomitant with dox-induction (Fig 6A-C & Suppl Fig 6E&G), demonstrating that the A-repeat and flXIST RNA similarly rely on histone deacetylation to initiate gene silencing. However, an important difference is seen if deacetylase inhibition *follows* dox-induction (by several days). For genes repressed by the A-repeat RNA, TSA treatment results in re-appearance of transcription foci (Fig 6A-C & Suppl Fig 6D), indicating that *ongoing* HDAC recruitment/activity was required, defining a reversible “HDAC-dependent” state. In contrast, the gene silencing induced by flXIST is no longer reversed but has become “HDAC-independent”, indicating other modifications by flXIST RNA prevent re-acetylation (Fig 6A-C & Suppl Fig 6F).

**FIGURE 6.**
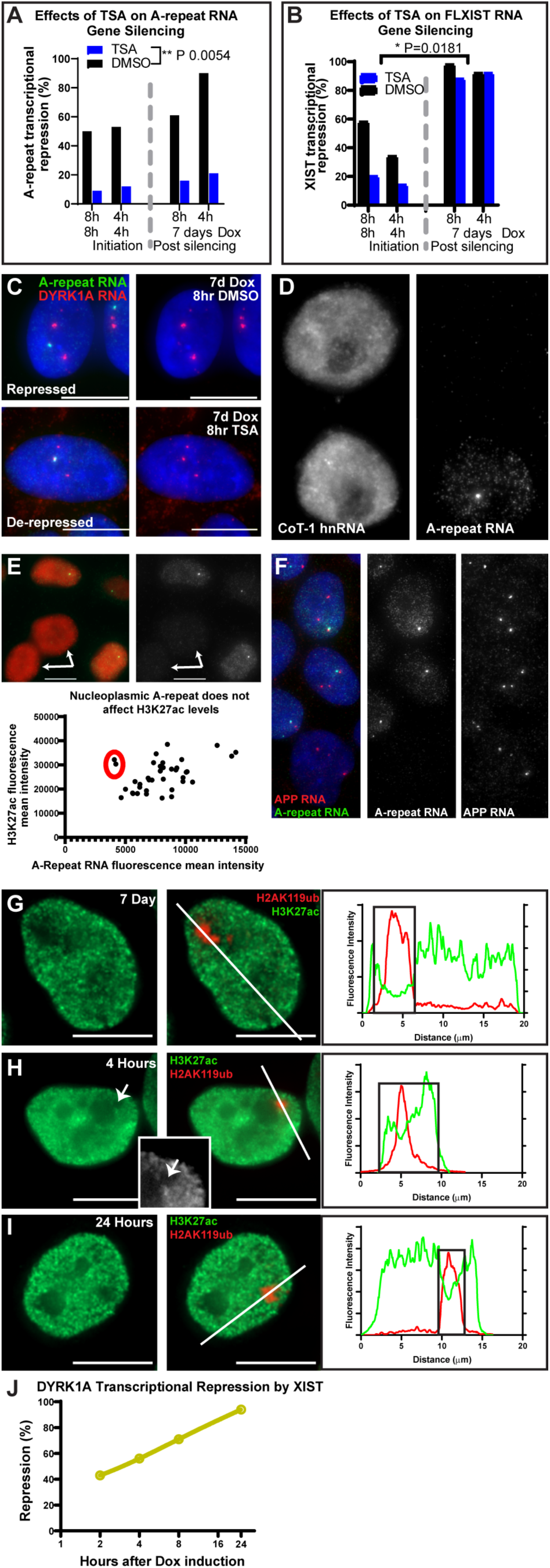
Histone de-acetylation essential for active gene silencing depends on high local density of A-repeat RNA. DAPI DNA is blue (C & F). **A-B)** Repression of DYRK1A transcription focus associated to A-repeat (A) or flXIST (B) by RNA FISH (Two-way ANOVA for significance). **C)** Representative FISH images quantified in A. Three color image (left) and green channel removed for clarity (right). (Suppl Fig 6 for more details). **D-F)** Analysis of whether substantial but lower levels of nucleoplasmic A-repeat RNA reduce histone acetylation or gene/hnRNA expression. **D)** CoT-1 RNA (left) in neighboring iPSCs induced and uninduced for A-repeat expression (right). **E)** H3K27ac (IF) and A-repeat RNA (FISH) in neighboring induced and uninduced iPSCs (green channel separated at right), with quantification of signal intensity (below), and cells lacking A-repeat RNA indicated (red circle: graph and arrows: images). **F)** APP and A-repeat RNA FISH. Red and green channels separated at right. **G-I)** H3K27ac and H2AK119ub (IF). Two-color images (right) and green channel alone (left). Linescans (far right) of two-color images (with white line), with edge of H2AK119ub signal indicated by black box in graphs. Close-up of green channel in black and white (H: insert) with H2AK119ub depletion indicated (arrows). **J)** Quantification of timing for silencing of DYRK1A gene, very near full-length XIST RNA transcription site.

**FIGURE 7:**
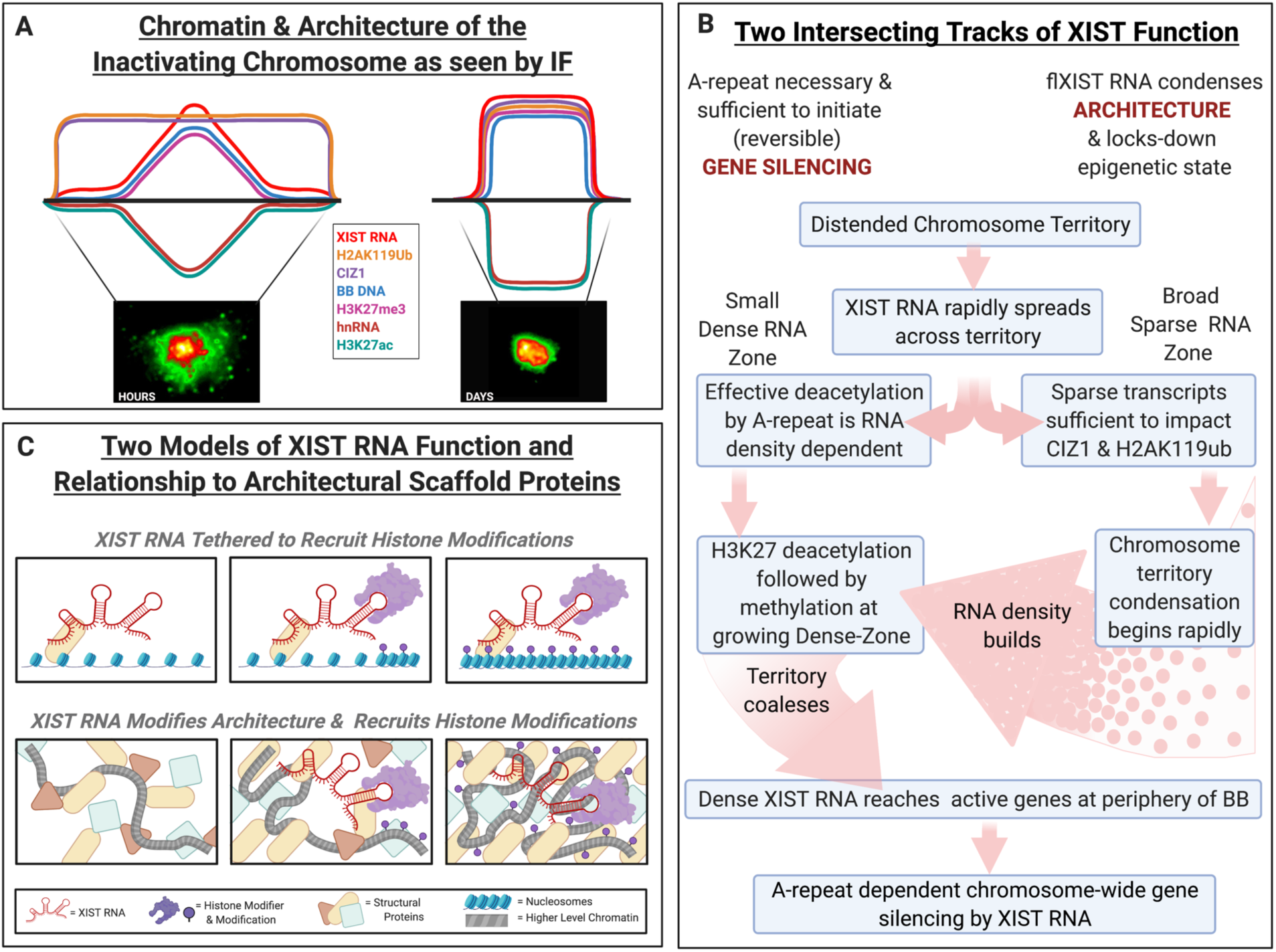
Models of XIST RNA function based on collective results. **A) Idealized composite linescan** modeling comparative distributions of biochemical and architectural changes on an individual inactivating chromosome territory, as observed *in situ*. *Left*, early hours of chromosome silencing process and *right*, after the process is more complete. Early in the process, XIST RNA is very unevenly distributed, in a large sparse zone and smaller dense RNA zone. Some steps are density dependent (V-shaped profiles early in the process) and others are not (low flat profiles early). All are more evenly distributed over the condensed DNA/RNA territory later in the process. **B) Two intersecting tracks of XIST function:** As detailed in text, the flow chart incorporates findings that support new concepts of XIST “two track” functions. Sparse XIST transcripts quickly act on architecture to condense the distended chromosome and coalesce a concentrated RNA territory, thereby facilitating density-limited function of the A-repeat required to silence active genes. **C) Two Models of XIST RNA Function and Relationship to Architectural Scaffold Proteins**. *<u>Traditional</u> <u>model (top):</u>* Non-chromatin matrix proteins simply tether XIST RNA to the chromosome in order to localize XIST RNA function, which is to recruit histone modifying enzyme(s) and trigger a histone modification cascade that silences genes and condenses the chromosome. *<u>Alternative model (bottom):</u>* XIST RNA acts early to modify architectural factors and also recruits histone modifiers. In this model XIST RNA is embedded with non-histone scaffold proteins, and acts directly to alter their arrangement to condense the large chromosome territory, and at the same time XIST RNA recruits enzymes for local histone modifications, some of which are required for gene silencing and further contribute to architectural condensation.

Unlike more stable “epigenetic” changes, histone deacetylation is known to have a broad role in transcriptional modulation that involves an ongoing dynamic balance between deacetylation (HDAC) and acetylation (HAT) (Pasini et al., 2010; Seto and Yoshida, 2014). Hence, efficient transcriptional repression by A-repeat RNA may require HDAC density sufficient to compete with HAT activity in transcriptionally active chromatin, in order to shift the balance towards repression. As indicated above, in addition to dense RNA foci, many cells contain substantial levels of A-repeat RNA throughout the nucleoplasm. To determine if these less concentrated A-repeat transcripts have any detectable impact on transcription we examined H3K27ac and hnRNA levels, as well as specific gene transcription foci in these cells, in direct comparison to neighboring cells with no A-repeat transgene expression. Cells with substantial nucleoplasmic A-repeat RNA showed no reduction in hnRNA (as detected by CoT-1 RNA)(Fig 6D) nor any change in H3K27ac levels (Fig 6E) compared to neighboring control cells lacking repeat-A RNA expression. Similarly, transcription foci for all genes studied above were only repressed when *in cis* with the dense A-repeat RNA foci, with no difference in signal presence or intensity for alleles within nucleoplasm containing substantial A-repeat RNA compared to adjacent non-expressing cells (e.g. Fig 6F).

The above results suggest that early in the inactivation process effective deacetylation by A-repeat sequences (within full-length XIST RNA) may be limited by the low RNA density across much of the chromosome; therefore we examined H3K27ac staining over the time-course on individual chromosomes, directly compared to staining for H2AK119ub (which also serves as a proxy for XIST RNA). In cells expressing flXIST RNA for seven days, when the process is essentially complete, a clearly defined dark “acetylation void” is apparent over the whole inactivated chromosome, also marked by H2AK119ub enrichment (Fig 6G). Whereas when H2AK119ub labels a broad territory detected at two hours, any chromosome-wide decrease in histone acetylation is barely discernible by IF (e.g. Fig 6H), and becomes more clearly evidenced only at ∼1 day (Fig 6I), although not to the extent readily apparent at Day 7. However, in some cells at early hours a dip in acetylation staining is apparent in the smaller dense zone (Fig 6H: insert); consistent with that, we tested and found that the DYRK1A gene, which occupies a unique location at the epicenter of XIST expression, is silenced more rapidly than genes tested above (Fig 6J). Collectively, these results support that early in the process histone deacetylation, sufficient to repress actively transcribed genes, is limited by the sparse nature of XIST transcripts across much of the distended territory; H3K27 deacetylation becomes more comprehensive only once the dense XIST RNA territory coalesces on a more compact chromosome territory. In contrast, changes to H2A and CIZ1 are rapidly widespread and much less limited by RNA density. Results here also demonstrate that the HDAC activity of the A-repeat element is necessary and sufficient to initiate gene silencing, and this can occur rapidly, but only if local RNA density is high. As discussed below, these results suggest a mechanistic explanation for a central point demonstrated here, that human XIST RNA triggers broad changes to territory architecture well before wide-spread silencing of most active genes.

## DISCUSSION

Many studies have described chromosome silencing in mouse ES cell models, but here use of an inducible-XIST human iPS cell system made it possible to define temporal steps in *human* chromosome silencing, in a pluripotent cell that fully supports XIST RNA function. The temporal order of major histone changes triggered by human XIST RNA (Fig1 and summarized in diagram in Fig 4), notably includes demonstration that H2AK119ub modification is seen before H3K27me3 in *human* chromosome silencing, which supports recent studies on this debated point in mouse chromosome silencing (Brockdorff, 2017). Moreover, our study not only defines timing but the spatial distribution of XIST RNA relative to various modifications in direct context of nuclear chromosome geography. Results support a new paradigm that initial differences in XIST transcript density are an important determinant of when and where specific modifications are triggered, as further discussed below. For reference, Fig 7A provides an idealized composite of how various chromatin changes, as observed *in situ*, corresponded to XIST RNA density on the chromosome territory. Our study then goes further to provide significant mechanistic insight into why XIST density may be more limiting to some steps, including active gene silencing. In addition to studying full-length human XIST RNA function, our study shows for the first time (in either species) that the 450 bp A-repeat RNA *alone* can shut-off transcription of endogenous genes in a local chromosome region. This discussion will be focused on key findings that support a unifying concept of these numerous findings. As outlined in Fig 7B, collective results support the novel concept that XIST RNA functions on two separable but inter-related tracks, one which modifies overall chromosome architecture, and one which silences canonical gene transcription. Results indicate that those two broad tracks of XIST function are controlled to some degree by separable domains of the long XIST transcript, with much of the long XIST transcript important to spread RNA across the chromosome, modify architecture, and build RNA density.

A key point that supports the overall concept is the discovery that barely detectable XIST transcripts rapidly spread across the very decondensed territory of pluripotent cells, creating a “sparse zone” which extends well beyond the initially small “dense zone” near the XIST locus. Detection of this very sparse RNA requires analysis at early time points as well as optimal RNA preservation and hybridization, and will be missed if imaging is dominated by the bright dense zone signal. These sparse transcripts indeed have a functional relationship to chromatin, as they rapidly trigger robust H2AK119ub and CIZ1 staining. Significantly, in contrast to bright H2AK119ub and CIZ1 across both zones, H3K27me3 is enriched in the dense zone, but not the sparse zone. This initially puzzling observation provided the first indication that specific aspects of XIST RNA function are more limited by RNA density than others. This is most apparent within a few hours of XIST induction, because as chromosome condensation proceeds for 1-2 days, the distended sparse zone coalesces into the larger more uniformly dense XIST RNA territory, as familiar in most studies.

Hence a dramatic change to overall architecture has occurred by Day 1 and 2, with DNA condensation coincident with a region lacking CoT-1 RNA; yet, silencing of several genes that map broadly across the chromosome did not occur until Day 3. This modification of architecture before gene silencing is reinforced by an earlier study of female mouse X-inactivation, in which several genes examined were similarly silenced after formation of the domain lacking CoT-1 RNA and RNA pol II, as well as studies finding this domain can form in ΔA-repeat mutants lacking gene silencing capability (Chaumeil *et al*., 2006). Insight into why genes are silenced days after sparse XIST RNA induces H2AK119ub and CIZ1 comes from analysis here of A-repeat RNA and histone deacetylation. Although the A-repeat alone can silence genes and flXIST (containing A-repeat) spreads rapidly, several findings support that A-repeat-mediated deacetylation requires higher density. Only concentrated foci of A-repeat (in minigene or flXIST) effectively reduce acetylation sufficiently to repress ongoing transcription of active genes. In keeping with this, it is known that regulation of many genes involves a dynamic balance of HAT and HDAC activity at a given site (Pasini *et al*., 2010; Seto and Yoshida, 2014). Given this ongoing balance, the A-repeat does not itself “silence” genes in the epigenetic sense, but represses activity as a reversible first step, which other XIST domains may function to epigenetically lock-down in subsequent steps. For example, deacetylation of H3K27 may be required for methylation of that same residue, which in turn blocks reacetylation. In sum, collective results support that effective histone deacetylation is required to initiate gene silencing, which is an A-repeat density-dependent step. Hence, condensation and coalescence build density of the XIST RNA territory, which then facilitates silencing of active genes throughout the chromosome. XIST RNA density would impact later steps as well, such as the formation of a highly stable, heterochromatic body through multivalent protein interactions, as recently described (Pandya-Jones *et al*., 2020). Here, results point to the impact of early RNA *in situ* distribution on specific chromatin modifications, including a density-limited first step required to initiate gene silencing.

Findings here highlight that XIST RNA function must be understood in the structural context of a nuclear chromosome territory that is overwhelmingly non-coding DNA, in which active genes are non-randomly organized. The natural XIST/Xist locus is in a repeat-rich, gene-poor region (which likely facilitates condensation of the Barr body), but our results show that if genes are very close to the XIST locus they can be repressed quickly, hence silencing of some genes or low-level lncRNAs likely occurs sooner. However, the geography of genes within the chromosome territory is not simply related to linear chromosome position, and four genes randomly distributed on the chromosome took days to silence (even though XIST induces broad changes within hours). In general genes localize disproportionately in the “outer reaches” of active chromosome territories, where many or most active protein coding genes cluster with inter-chromosomal SC-35 speckles (Chen and Belmont, 2019; Hall and Lawrence, 2010; Kurz *et al*., 1996; Shah et al., 2018; Shopland et al., 2003; Smith *et al*., 2020; Xing *et al*., 1993). Moreover, in the distended chromosome of pluripotent cells most active genes begin far from the initially small dense XIST RNA zone and remain active until condensation draws them towards the condensed XIST RNA territory, where they remain in fully silenced X-chromosomes (Clemson *et al*., 2006; Hall and Lawrence, 2010); similar inward movement of distant genes to the periphery of the territory is evident during mouse X-inactivation (Chaumeil *et al*., 2006).

Our analysis of very early biochemical changes juxtaposed with changing chromosome architecture of single cells reveals findings difficult to discern by extraction-based analyses of averaged populations. However, the latter provide valuable information on sequence context; most notably, the observations that XIST RNA first modifies H2AK119ub and H3K27me3 in intergenic sequences and less active regions (before active genes) (Nesterova *et al*., 2019; Zylicz *et al*., 2019) fits well with our conclusion that XIST RNA first modifies the architecture of the largely non-coding chromosome, and later silences active genes in regions requiring high RNA density for histone deacetylation. Regions that are non-genic or lowly expressed may be more readily impacted by sparse XIST transcripts and contribute to early DNA condensation of the Barr body devoid of Cot-1 hnRNA.

Given this evidence that condensation is not just a by-product but an early facilitator of broad gene silencing, then more focus is warranted on how XIST modifies cytological-scale structure. Findings here suggest a new hypothesis, as modeled in Fig 7D. We show that sparse XIST transcripts immediately impact CIZ1, a nuclear matrix protein. Current models suggest matrix proteins such as SAF-A and CIZ1 serve to tether XIST RNA to the chromosome to localize XIST’s function via the histone cascade (Fig 7C). Instead, we propose that CIZ1 is actually an early *target* of XIST RNA’s “second-track” of function, to directly modify large-scale architecture (Fig 7D). In this model, both XIST RNA and CIZ1 are integral components of a complex chromosome territory scaffold. Whether XIST RNA binding modifies CIZ1 conformation (to reveal the epitope) or recruits CIZ1, it is likely that XIST RNA is not just anchored by CIZ1 but impacts its interactions with other components. Given that CIZ1 binds DNA and becomes detected by IF after chromatin removal (Swarts *et al*., 2018), an attractive hypothesis is that XIST RNA binding changes CIZ1-DNA interaction. A recent study from our lab shows that long euchromatin-associated CoT-1 hnRNAs (lncRNAs and pre-mRNAs) platform an insoluble RNP scaffold which counters chromosome condensation (Creamer *et al*., 2021) hence this recent study and other evidence indicates XIST RNA may displace or silence RNAs that promote open euchromatin (Hall et al., 2014). Early in the process sparse XIST RNA may release chromatin in the distended territory from euchromatin scaffold factors. However, condensation would likely also be promoted by coalescence of dense XIST transcripts with other factors near the epicenter, the XIST locus. In fact, a recent study describes density-dependent phase separation of multivalent proteins with XIST RNA, which is suggested to further lock-down an XIST RNA-independent heterochromatic state after cell differentiation (Pandya-Jones *et al*., 2020). Results here reveal that initially there are sharp regional differences of XIST RNA density across larger chromosome territory, which in turn has unanticipated importance for understanding early and distinct steps of XIST RNA function.

Finally, work on the XIST A-repeat was motivated in part by our goal to advance “translational epigenetics” with XIST. A major impact of this work is the enhanced prospect that small XIST derivatives can be developed translationally for the common, unaddressed problem of chromosomal duplication disorders. While this study focuses on the fundamental biology, findings here provide the basis for work of high translational potential, to be amplified elsewhere.

## ACKNOWLEDGMENTS

We appreciate the support of the NIH—grants R01R35GM122597, R01HD091357, and R01HD094788 to J.B.L. and F32 AG056131-01 to MV. We thank members of the Lawrence lab for support in various ways, including thoughtful discussions and critical analysis of this study. We appreciate Jennifer Moon providing advance access to her RNA seq data and analysis of CIZ1 expression before and after differentiation.

---

**SUPPL FIG 1:**
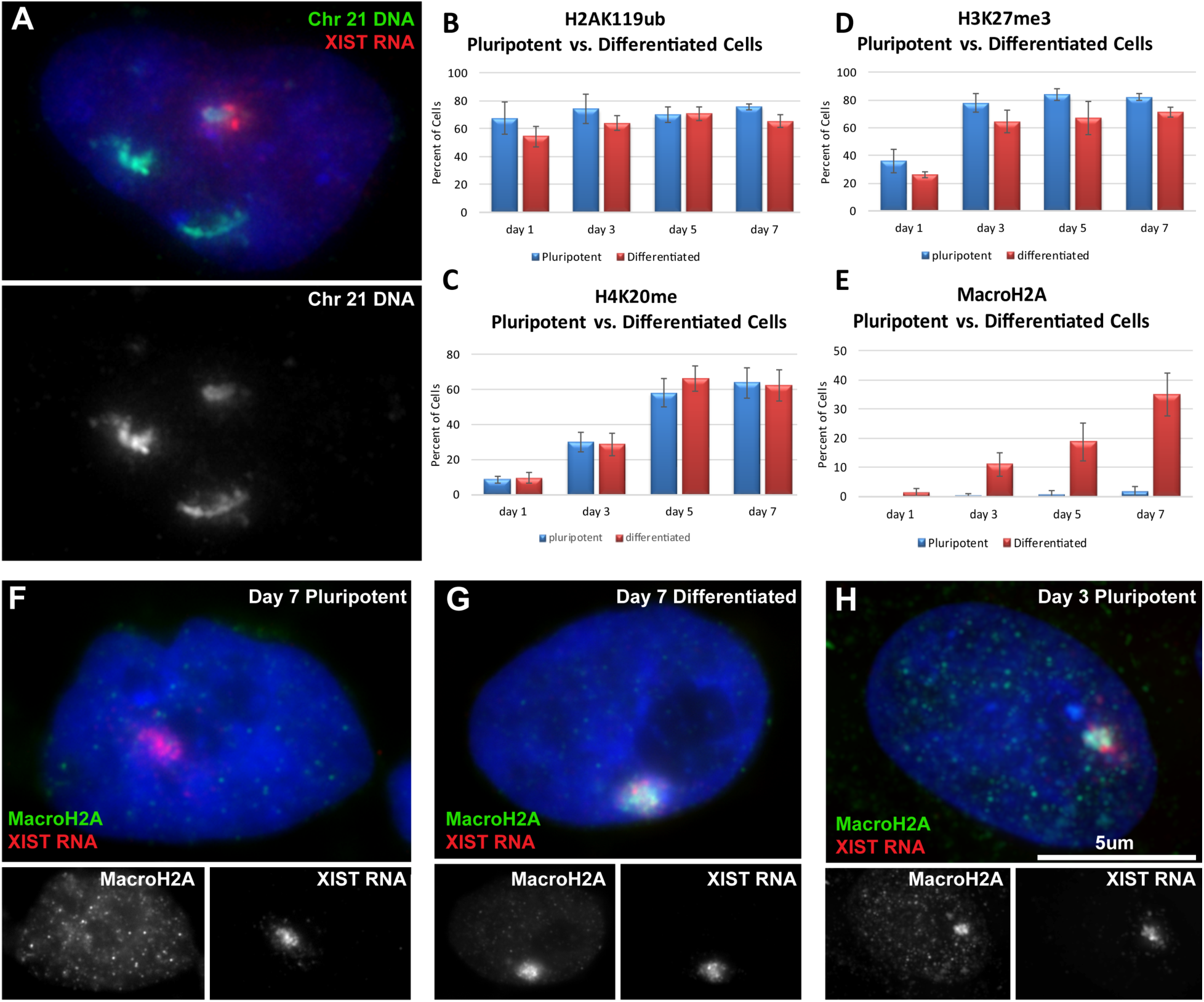
XIST RNA compacts an initially distended chromosome and heterochromatic hallmarks are largely similar between pluripotent and differentiated cells. DAPI DNA is blue (A, F-H). **A)** Chr21 library DNA in Down syndrome iPSC showing 3-chr21, with XIST RNA FISH indicating compacted transgenic chromosome. Green channel separated below. **B-E)** The timing of chromatin hallmarks scored in pluripotent and differentiating iPSCs during 7 days of XIST expression. Only macroH2A (E) shows a significant difference between differentiating and pluripotent cultures. **F-H)** MacroH2A enrichment was only observed *upon differentiation* in human iPSCs (F & G) and ES cells(Hoffman *et al*., 2005) under older growth and maintenance protocols using inactivated feeders. However, using modern iPSC feeder-free culture conditions we observe macroH2A enrichment beginning on day 3 in *pluripotent cells* (H), suggesting modern culture methods may change epigenetic plasticity of these cells. Red and green channels separated below main images.

**SUPPL FIG 2:**
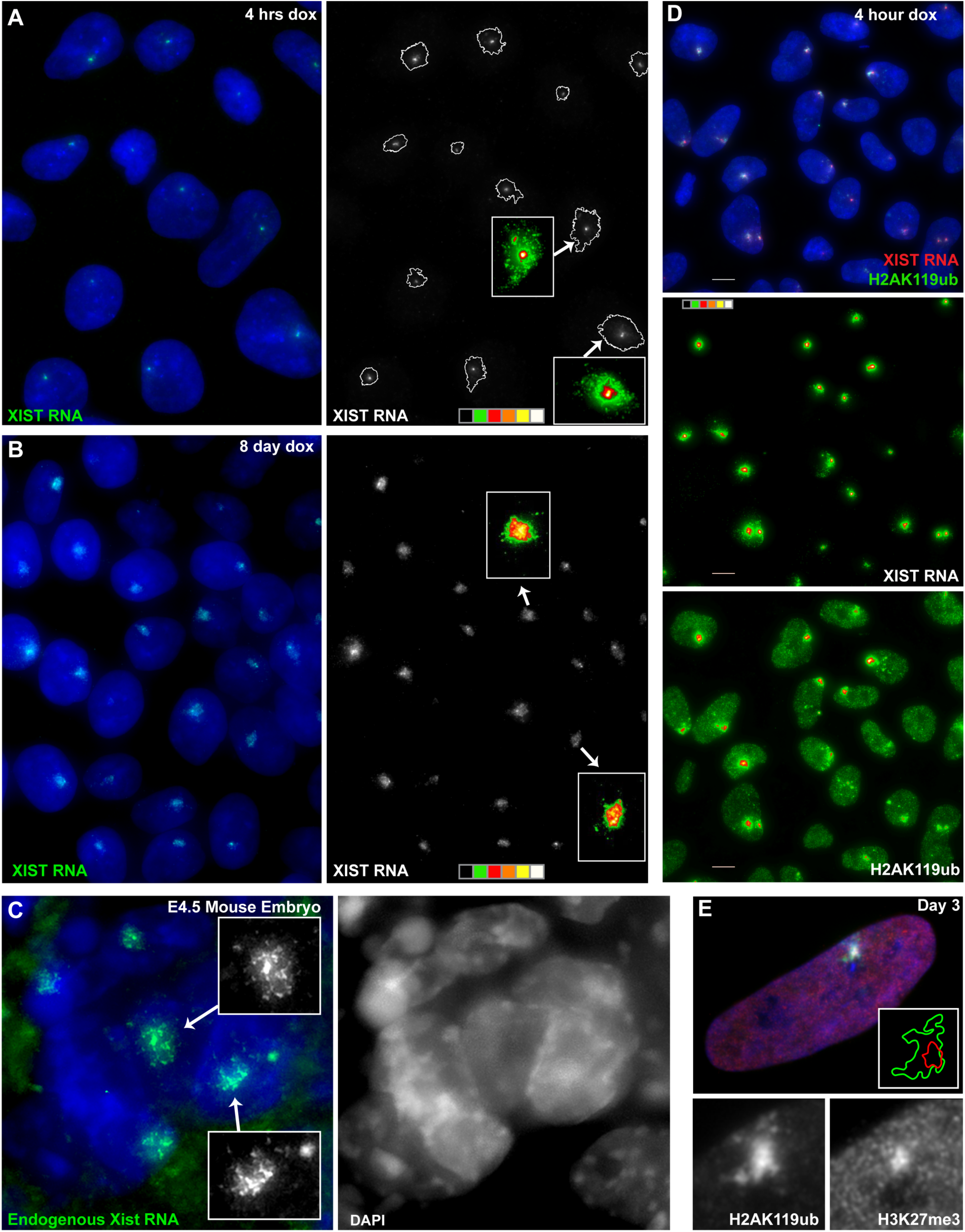
Low level spread of XIST RNA is seen early in the process and may often be missed but they impact chromatin. DAPI DNA is blue (all images). **A-B)** A field of iPSC at 4 hours (A) and 8 days (B) show the change in the XIST RNA territory over time. The green channel is separated at right with threshold edges of the 4-hour XIST territory outlined. Inserts show two representative XIST RNA signals (arrows) with a 6-color heat map of pixel intensity showing sparse (green) and dense (red-white) zones. Note: Fig 2 in main text shows region of same 4hr field. **C)** During X-inactivation in very early mouse embryos, endogenous Xist-RNA also exhibits a large sparse dispersal rather than a small compact cloud (surrounding trophectoderm cells are not included in the image). The green channel is separated for select Xist RNA territories in inserts. Blue channel for entire cell mass separated at right. **D)** Field of 4hr induced cells with H2AK119ub (IF) enrichment under XIST RNA (FISH) territories. Note, not all cells in this field responded to induction and expressed XIST. A 6-color heat map of XIST RNA and H2AK119ub IF pixel intensity is separated below. **E)** H2AK1129Ub enrichment is seen across the entire XIST RNA sparse zone, while H3K27me3 is only seen over the center dense XIST RNA zone. Red and green channels separated below, and illustration of the threshold-edge of the signals in insert.

**SUPPL FIG 3:**
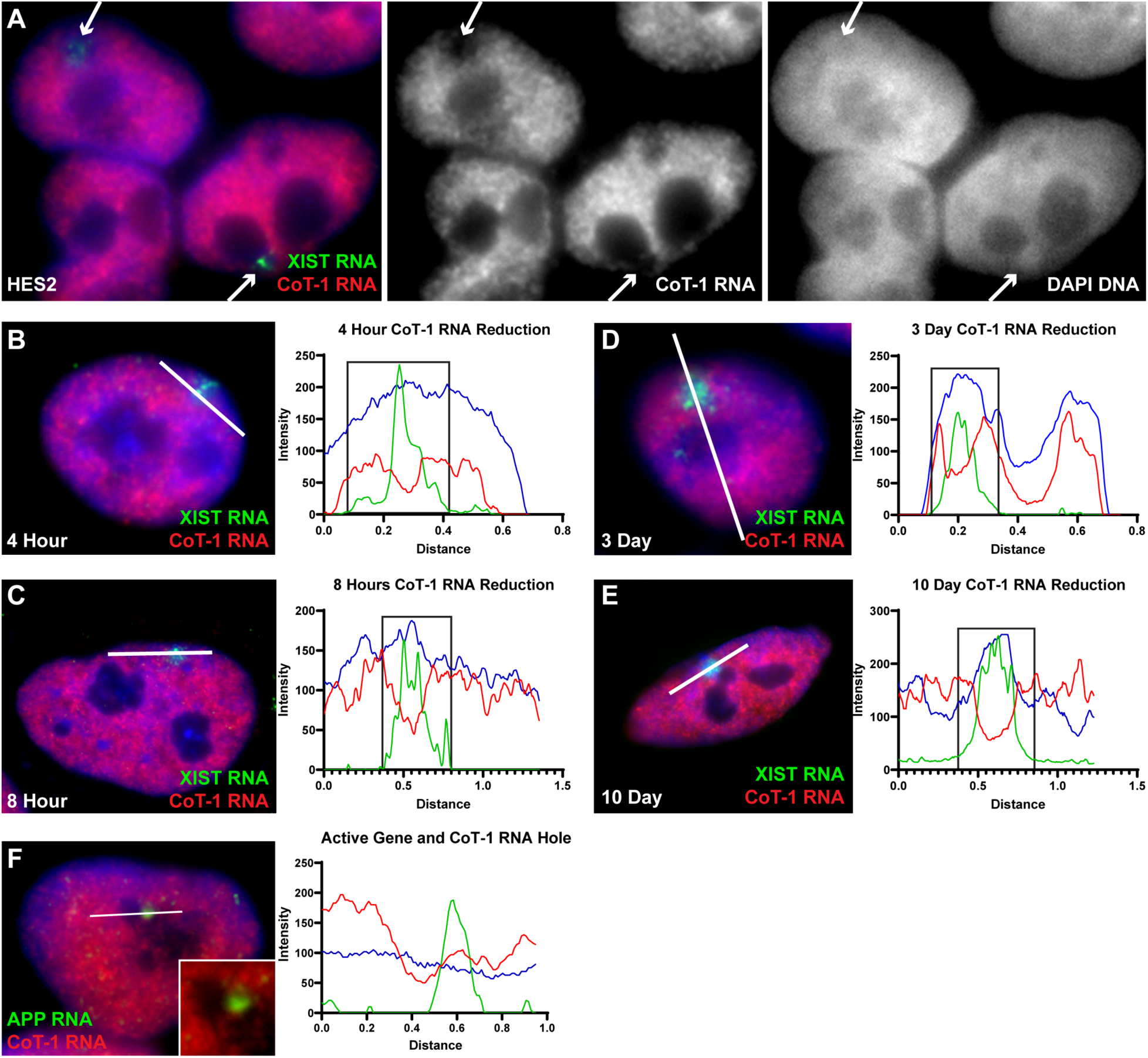
CoT-1 RNA “hole”/Barr body formation over the inactivating chromosome. DAPI DNA is blue (all images). **A)** DAPI dense Barr bodies (BB) are not easy to detect in all cell preps (particularly pluripotent cells), making the presence of a CoT-1 RNA hole the most reliable way of detecting the BB. Red and blue channels separated at right with location of inactive chromosome (with XIST RNA expression) indicated (arrows). **B-E)** Reduction of CoT-1 RNA over XIST RNA territory in 4hr, 8hr, 3-day and 10-day nuclei. Linescans across regions delineated in 3-color images (white lines) are at right. Edges of XIST RNA territory indicated by black boxes in graphs. F) APP and CoT-1 RNA FISH show APP transcription focus at edge of CoT-1 RNA hole prior to silencing in iPSC. Linescan across region (white line) at right. Closeup of region, with Blue channel removed, in insert.

**SUPPL FIG 4:**
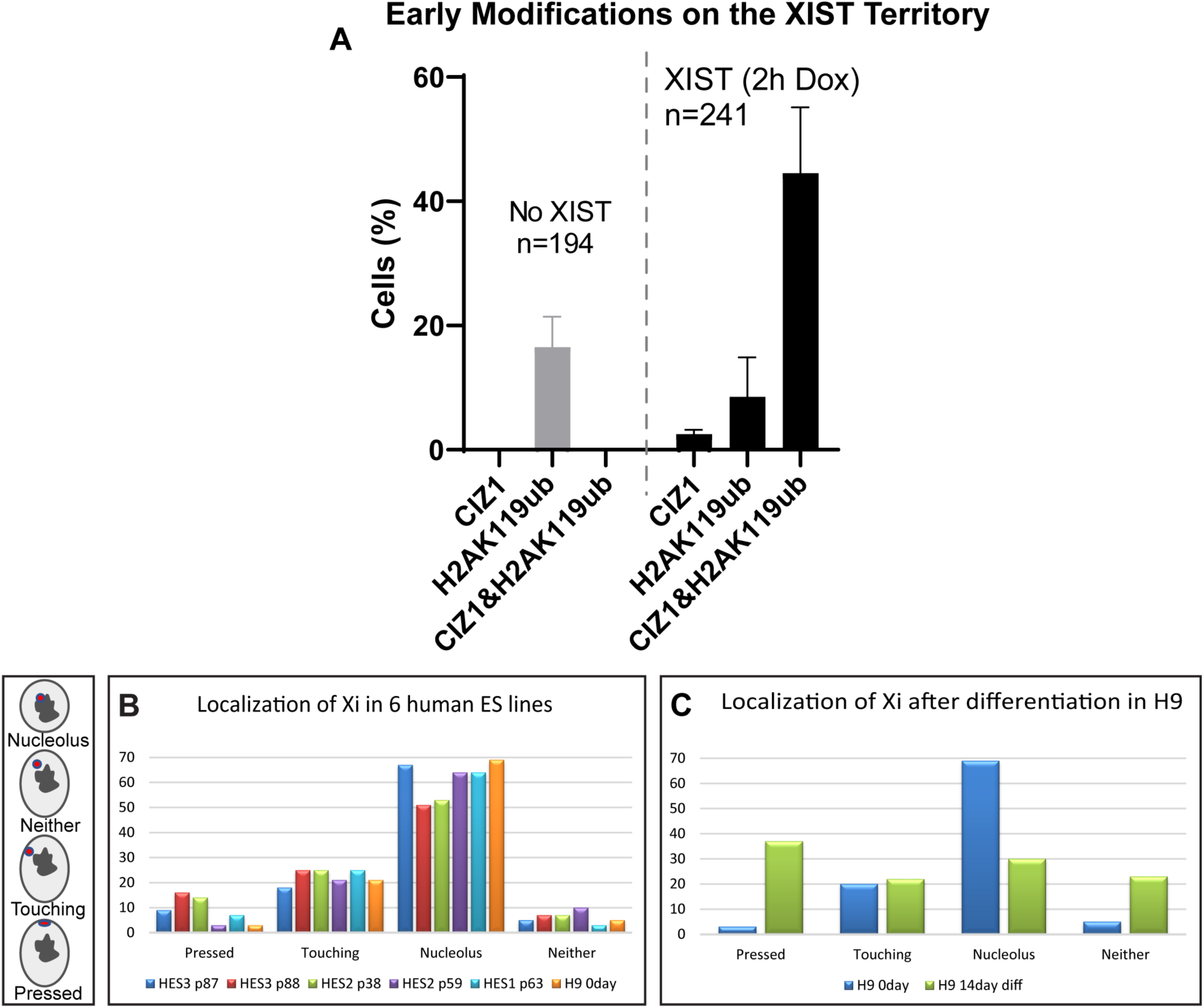
XIST RNA impacts the scaffold early but chromosomal movement to nuclear periphery is late and requires differentiation. **A)** Detection of CIZ1 and H2Ak119ub accumulation before and after XIST expression. To determine whether CIZ1 or H2Ak119ub appears first, simultaneous staining for both proteins was done at 2hrs of XIST induction (RNA hybridization was not included to optimize detection of both antibodies). In most cells, both proteins were detected and co-localized in a single bright cloud, presumed to be the XIST transcription focus. But a small fraction of cells at 2 hours contained an enriched focus of just one signal (CIZ1 or H2AK119ub). Because some non-induced cells already contained H2AK119ub foci, we can conclude that a small subset of cells are enriched for CIZ1 without H2AK119ub modification, but we cannot determine if the reverse is also true. **B-C)** Nuclear location of the precociously inactivated Xi in several *pluripotent* human ES cell lines (B) and in the H9 hESC line *after differentiation* (C). Illustrations of each type of chromosome locations scored is at left.

**SUPPL FIG 5:**
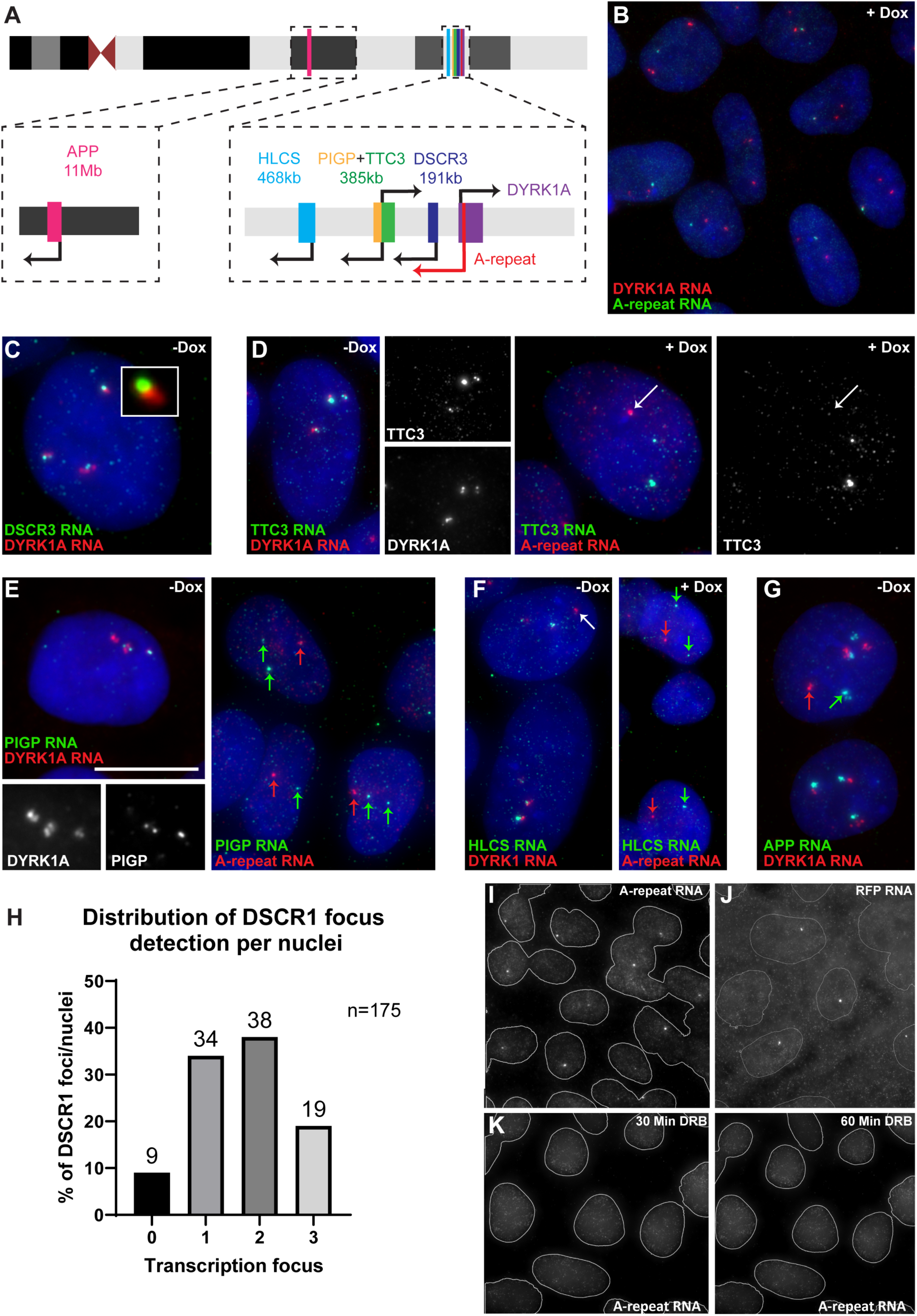
High density focal A-repeat RNA silences nearby genes while low levels of A-repeat RNA distribute broadly but remain in the nucleus. DAPI DNA is blue (B-G). **A)** Diagram of Chr21 gene loci examined for silencing by A-repeat RNA. **B)** Field of induced cells with DYRK1A and A-repeat RNA FISH. **C)** DSCR3 (also TTC3 (D), PIGP (E), HLCS (F) & APP (G)) RNA foci were scored in relation to DYRK1A RNA foci to ascertain hybridization frequency in uninduced cells. These were then compared to induced samples to determine silencing frequency by A-repeat. **D)** TTC3 & DYRK1 RNA FISH in uninduced cells (left) and TCC3 & A-repeat RNA FISH in induced cells (right). Separated channels in black & white as indicated. Silenced allele indicated (arrow). **E)** PIGP & DYRK1 RNA FISH in uninduced cells (left) and PIGP & A-repeat RNA FISH in induced cells (right). Separated channels in black & white as indicated. Silenced allele (red arrow) and expressed alleles (green arrows) indicated. **F)** HLCS & DYRK1 RNA FISH in uninduced cells (left) and HLCS & A-repeat RNA FISH in induced cells (right). Reduced hybridization efficiency resulted in some DYRK1 foci not having a corresponding HLCS focus (white arrow). Silenced allele (red arrow) and expressed alleles (green arrows) also indicated. **G)** Because the APP gene is 11MB away from the DRYRK1 locus (where the A-repeat is targeted), they can be far apart in some nuclei (red/green arrows), but three foci were apparent in all cells whether induced or uninduced. **H)** Example illustrating genes examined were not monoallelically expressed; even for TFs that did not produce strong *in situ* nuclear signals, we still detect two or three expressing alleles in many cells. This point is significant because it argues against single-cell seq analysis interpreted to show that most genes express from just one allele, even in trisomic cells (Stamoulis et al., 2019). **I-J)** A single channel image of A-repeat and RFP RNA (FISH) I) a field of cells shows a bright RNA focus and a lower density dispersed nucleoplasmic RNA signal filling individual nuclei. A-repeat RNA is restricted to the cytoplasm, while RFP mRNA is transported to the cytoplasm for translation. Nuclei outlined in white. **K)** A-repeat transcription foci are gone after 30min of transcriptional inhibition (left) leaving only dispersed signal delineating nuclei, and another 30min is required for complete loss of signal.

**SUPPL FIG 6:**
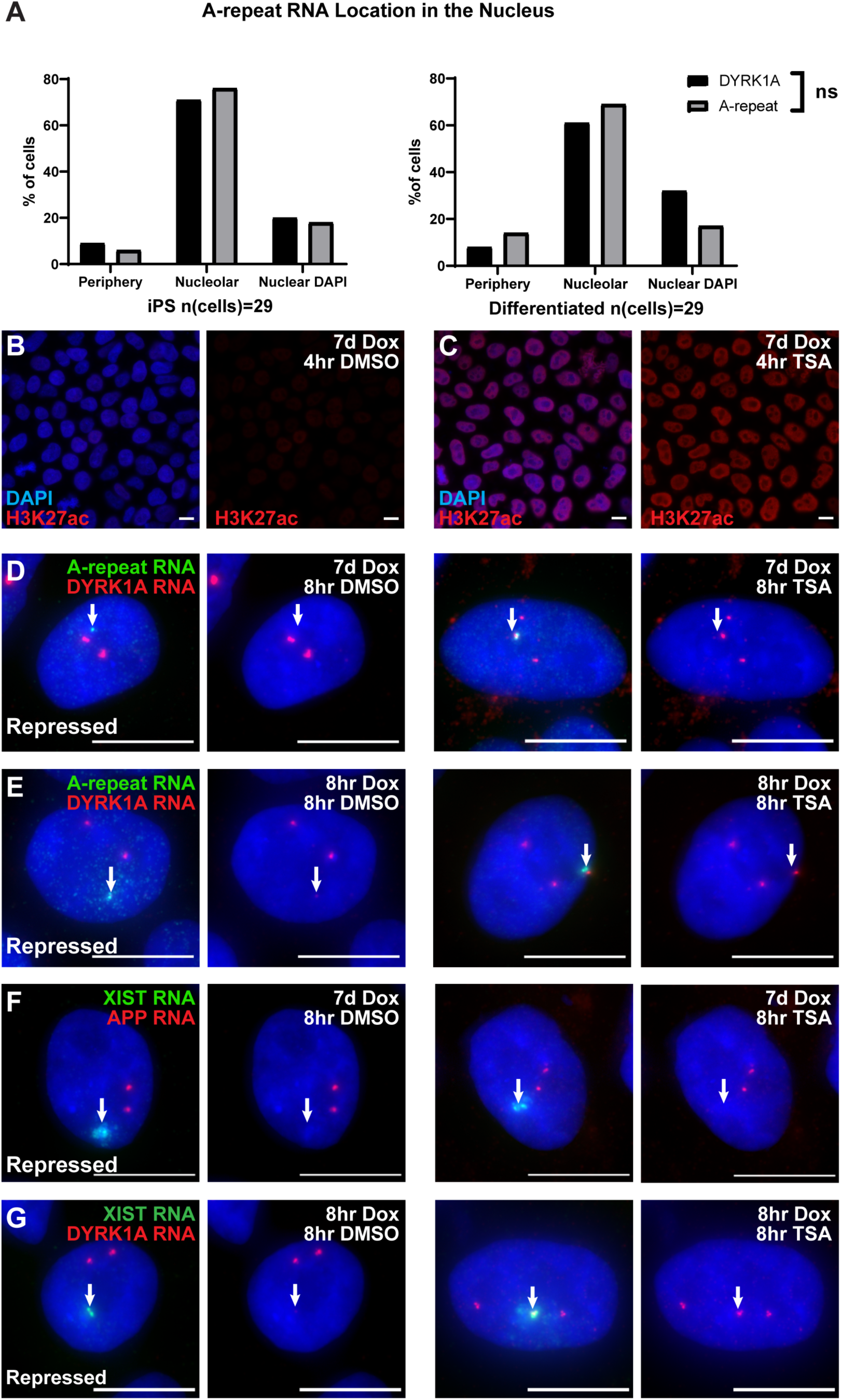
Nuclear periphery is not involved in gene silencing but TSA treatment during silencing reveals an HDAC-dependent and HDAC-independent silencing state. DAPI DNA is blue (B-G). **A)** Nuclear localization of A-repeat RNA focus (on transgenic Chr21) compared to DYRK1A alleles on non-transgenic Chr21s in pluripotent and endothelial differentiated iPSCs. **B-C)** H3K27ac (IF) in cells treated with TSA or DMSO for 4 hours. **D-E)** Representative example of A-repeat and DYRK1A RNA FISH images used in Fig 6A quantification. TSA treatment (or DMSO alone) *following* gene silencing (D) or *during* gene silencing (E). **F-G)** Representative example of flXIST and DYRK1A/APP RNA FISH images used in Fig 6B quantification. TSA treatment (or DMSO alone) *following* gene silencing (F) or *during* gene silencing (G). DYRK1A was used for short-term TSA treatment during flXIST mediated chromosome silencing, since APP took days to silence.

## STAR★Methods

### LEAD CONTACT AND MATERIALS AVAILABILITY

Further information and requests for resources and reagents should be directed to and will be fulfilled by the Lead Contact, Jeanne Lawrence.

### EXPERIMENTAL MODEL AND SUBJECT DETAILS

#### Human Cells

This study was mainly performed in iPSC derived from a male with Down syndrome kindly provided by G.Q. Daley (Park et al., 2008). XIST-transgenic lines were accomplished and characterized in Jiang et al. (2013). A-repeat transgenic lines were accomplished for this study as described below. To illustrate certain points, other cell lines were used such as H9 hESC and female TIG-1 (normal human lung primary fibroblast).

iPSCs and ESC were maintained on irradiated mouse embryonic fibroblasts (iMEFs) (R&D Systems, PSC001) in hiPSC medium containing DMEM/F12 supplemented with 20% Knockout Serum Replacement, 1mM glutamine, 100mM non-essential amino acids, 100mM b-mercaptoethanol and 10ng/ml FGF-β at 37°C with 20% O_2_ and 5% CO_2_. Cultures were passaged every 5–7 days with 1mg/ml of collagenase type IV. In later studies, cells were grown in Essential 8 medium on plates coated in vitronectin 0.5 ug/cm^2^. Cells were passaged at 80% confluency every 3-4 days with 0.5mM EDTA. TIG-1 line was cultured in MEM 15% FBS. Cells were periodically tested for mycoplasma.

Expression of XIST and the A-repeat was induced with doxycycline (500ng/ml) while maintained as pluripotent, or directly upon differentiation. Random differentiation was achieved by removing iPS cells from feeder layer and feeding with DMEM/F12, 4% Knockout Serum Replacement, 100mM Non-essential amino acids, 1mM L-glutamine, 100mM β-mercaptoethanol. iPS cells were differentiated into endothelial cells with Gsk3 inhibitor (as in (Bao et al., 2016) and Moon, submitted) in LaSR basal media (formulated from Bao 2016 (Bao *et al*., 2016)) with 6μM CHIR99021 for the first two days. Endothelial precursor cells were purified using a CD34 MicroBead Kit (Miltenyi Biotec, cat# 130-100-453); and maintained in EGM2 (Lonza, cat# CC-3162) (with 5μM Y-27632 for the first day) on vitronectin coated plates. NPC differentiation was performed as (Czerminski and Lawrence, 2020).

For transcriptional studies, HDAC and protein phosphatase 1 inhibition, cells on coverslips were incubated with 50ug/ul 5,6-Dichlorobenzimidazole 1-β-D-ribofuranoside (DRB), 5-10uM trichostatin-A (TSA) and 3uM Tautomycin for the indicated time. Cells were then fixed as indicated below for RNA FISH.

#### Mouse Cells

Male mouse J1 ES cells containing a doxycycline-inducible Xist cDNA transgene integrated on Chr-11 (clone #65)(Wutz *et al*., 2002) was also briefly used to illustrate some points. These cells were maintained in DMEM (GIBCO), 15% fetal calf serum (FCS, Hyclone), on mitomycin inactivated (10ug/ml mitomycin C for 2 hours at 37C) STO fibroblast feeder cells (SNL76) that produce LIF from an ectopic transgene. mES cells were differentiated by removing colonies from feeders (through two, two-hour sequential separations of single cell suspension onto gelatinized flasks) and distributing them as a single cell monolayer on gelatinized (0.1% porcine skin gelatin) flasks in the presence of 100nM all-*trans*-retinoic acid. Xist RNA expression was induced with 1 ug/ml doxycycline at the same time. Time points were taken by trypsinizing the cells and plating them as a monolayer onto coverslips coated with CellTak (BD) (following manufacturers protocol) for 1 hour before fixation.

### METHOD DETAILS

#### pTRE3G-A-Repeat-EF1a-RFP::DYRK1A plasmid

A-Repeat, and backbone with arms to DYRK1A, were PCR amplified from pTRE3G-XIST(Jiang *et al*., 2013). The EF1αRFP was amplified from plasmid HR700PA-RFP (System Biosciences). The five PCR products were GIBSON assembled. Primer sequences are listed in table.

#### Inducible A-repeat cell line

The inducible A-repeat transgene was targeted to the first intron of the DYRK1A locus in chromosome 21, the transactivators were targeted to chromosome 19 AAV site in Down syndrome iPS cells as described in (Jiang *et al*., 2013), but using PBAE (poly(β-amino ester), C320 (generously provided by the Anderson Lab, MIT(Eltoukhy et al., 2012; Zugates et al., 2007)). Briefly, Down syndrome iPS cell parental line provided by G. Q. Daley (Children’s Hospital Boston)(Park *et al*., 2008) were grown to exponential phase and cultured in 10mM of Rho-associated protein kinases (ROCK) inhibitor (Calbiochem; Y27632) 24h before transfection. A total of 55mg DNA including five plasmids (pTRE3G-A-Repeat-EF1a-RFP, DYRK1A ZFN1, DYRK1A ZFN2, rtTA/puro and AAVS1 ZFN) with 6:1 ratio of A-repeat:rtTA/puro were mixed with 1:20 ratio of PBAE Polymer and incubated with cells for four hours. Cells were washed with media and kept overnight with Essential 8 medium and rock inhibitor. Next day, cells were selected for puromycin resistance. Red clones were isolated. Expression of the A-repeat was induced with 500ug/ul doxycycline. Clones that lost the red fluorescence upon dox induction were used for this study. Expression of A-repeat was validated by RNA FISH and proper targeting by colocalization of the A-repeat and DYRK1A RNA transcription foci by RNA FISH.

RFP and DYRK1 RNA were usually detected in separate but adjacent transcription foci. However, we noticed that upon dox induction, some A-repeat transcripts also contained downstream sequences for RFP and DYRK1A in a colocalizing focus, but this co-localized RFP/DYRK1A signal was restricted to the A-repeat transcription focus, and appeared only in the presence of dox, suggesting read-through. Although this RFP/DYRK1 RNA read-though signal persisted in the presence of dox, the RFP protein was no longer present, indicating gene silencing. Thus, no functional mRNA for RFP or DYRK1A was expressed from this locus upon dox induction and gene silencing.

Cells kept in the presence of puromycin selection expressed the A-repeat transgene in almost 100% of cells. The frequency of cells expressing A-repeat dropped over time when grown in the absence of puromycin due to stochastic silencing of the tet-activator. These non-inducing cells were used as internal “non-expressing” controls for many experiments.

#### DNA and RNA FISH and immunostaining

These protocols were carried out as previously described (Byron et al., 2013; Clemson *et al*., 1996). Cells were fixed for RNA in situ hybridization as described in (Byron *et al*., 2013). Briefly, cells cultured on coverslips were extracted with triton X-100 for 3 min and fixed in 4% paraformaldehyde in phosphate-buffered saline (PBS) for 10 min. Cells were then dehydrated in 100% cold ethanol for 10 min and air-dried. Cells were then hybridized with biotin-11-dUTP or digoxigenin-16-dUTP (Dig) labeled nick translated DNA probes. DSCR3, TTC3, PIG3, HLCS DNA probes were obtained by amplifying ∼10Kb gene regions from the DS iPS genomic DNA and cloned into TOPO vector A, cold TOPO vector was added to the hybridization mixture of TOPO constructions to decrease background.

For hybridizations, 50ng of labeled probes with CoT-1 competitor were resuspended in 100% formaldehyde, followed by denaturation at 80°C for 10 min. Hybridizations were performed in 1:1 mixture of denatured probes and 50% formamide hybridization buffer supplemented with 2U/μl of RNasin Plus RNase inhibitor for 3 h or overnight at 37°C. Cells were then washed three times for 20 min each, followed by detection with anti-dig or streptavidin fluorescently conjugated secondary antibody. DNA was stained with DAPI. In simultaneous DNA/RNA FISH (interphase targeting assay), cellular DNA was denatured and hybridization was performed with 2U/ml of RNasin Plus RNase inhibitor to preserve RNA. For immunostaining with RNA FISH, cells were immunostained first with primary antibodies containing RNasin Plus and fixed in 4% paraformaldehyde after detection, before RNA FISH.

Most antibodies were diluted at 1:500 ratio. X chromosome was detected with whole chromosome paint probe (ID Labs Biotechnology), following manufacturers instructions.

### QUANTIFICATION AND STATISTICAL ANALYSIS

#### Image analysis

Cells were imaged on a Zeiss AxioObserver 7, equipped with a 100x Plan-Apochromat oil objective (NA 1.4) and Chroma multi-bandpass dichroic and emission filter sets (Brattleboro, VT), with a Flash 4.0 LT CMOS camera (Hamamatsu). Z stacks were taken for each field to evaluate detectable transcription foci. To evaluate if the A-repeat silenced nearby genes, we compared the frequency that a gene’s transcription focus was in close proximity to DYRK1A or RFP RNA foci in the absence of doxycycline, to the frequency in the presence of doxycycline. Images show a plane from the z stack or a MIP (indicated). Most experiments were carried out a minimum of 3 times, with typically 100-300 cells scored in each experiment. Key results were confirmed by at least two independent investigators. Line scans were done in Image J or ZEN 3.1 (Profile function) and plotted in Prism. Heat maps were created with Image J (fuji). Images were minimally enhanced for brightness and contrast to resemble what was seen by eye through the microscope. Due to the low intensity of the sparse XIST RNA in the sparse-zone and the dynamic range between that and the transcription focus, the initial sparse spread of XIST RNA may often be missed if cells are only observed on a computer screen (with poor dynamic range) rather than by eye under a microscope. Sparse-zone XIST can also be missed in images that are processed too much, or by super-resolution techniques that often reduce sensitivity.

Transcriptomic data was generated for a different study (Moon et al. submitted). Briefly, data was originated from 4 transgenic lines. NPC were achieved as in (Czerminski and Lawrence, 2020) (Czerminski, 2020) and collected for sequencing on diff day 14 (dox at diff day 0) while endothelial cells were differentiated with Gsk3 inhibitor as in (Bao *et al*., 2016) and collected for sequencing on diff day 12. RNA seq analysis was performed using EdgeR (McCarthy et al., 2012), using normalized cpm values. Figure uses log2 values.

